# Robust chronic convulsive seizures, high frequency oscillations, and human seizure onset patterns in an intrahippocampal kainic acid model in mice

**DOI:** 10.1101/2021.06.28.450253

**Authors:** Christos Panagiotis Lisgaras, Helen E. Scharfman

## Abstract

Intrahippocampal kainic acid (IHKA) has been widely implemented to simulate temporal lobe epilepsy (TLE), but evidence of robust seizures is usually limited. To resolve this problem, we slightly modified previous methods and show robust seizures are common and frequent in both male and female mice. We employed continuous wideband video-EEG monitoring from 4 recording sites to best demonstrate the seizures. We found many more convulsive seizures than most studies have reported. Mortality was low. Analysis of convulsive seizures at 2-4 and 10-12 wks post-IHKA showed a robust frequency (2-4 per day on average) and duration (typically 20-30 sec) at each time. Comparison of the two timepoints showed that seizure burden became more severe in approximately 50% of the animals. We show that almost all convulsive seizures could be characterized as either low-voltage fast or hypersynchronous onset seizures, which has not been reported in a mouse model of epilepsy and is important because these seizure types are found in humans. In addition, we report that high frequency oscillations (>250 Hz) occur, resembling findings from IHKA in rats and TLE patients. Pathology in the hippocampus at the site of IHKA injection was similar to mesial temporal lobe sclerosis and reduced contralaterally. In summary, our methods produce a model of TLE in mice with robust convulsive seizures, and there is variable progression. HFOs are robust also, and seizures have onset patterns and pathology like human TLE.

**SIGNIFICANCE STATEMENT:** Although the IHKA model has been widely used in mice for epilepsy research, there is variation in outcomes, with many studies showing few robust seizures long-term, especially convulsive seizures. We present an implementation of the IHKA model with frequent convulsive seizures that are robust, meaning they are >10 sec and associated with complex high frequency rhythmic activity recorded from 2 hippocampal and 2 cortical sites. Seizure onset patterns usually matched the low-voltage fast and hypersynchronous seizures in TLE. Importantly, there is low mortality, and both sexes can be used. We believe our results will advance the ability to use the IHKA model of TLE in mice. The results also have important implications for our understanding of HFOs, progression, and other topics of broad interest to the epilepsy research community. Finally, the results have implications for preclinical drug screening because seizure frequency increased in approximately half of the mice after a 6 wk interval, suggesting that the typical 2 wk period for monitoring seizure frequency is insufficient.

**HIGHLIGHTS:**

• Our implementation of the IHKA model led to robust chronic spontaneous convulsive seizures in mice
• Convulsive seizures were synchronized in both hippocampi and two cortical sites
• Seizure duration increased between 2-4 wks and 10-12 wks after IHKA
• Convulsive seizures fit LVF and HYP types found in human temporal lobe epilepsy
• HFOs (>250 Hz) were common, at >1 location, and were both ictal and interictal

## INTRODUCTION

The primary purpose of this study was to modify existing methods for a widely used animal model of temporal lobe epilepsy (TLE) so that robust chronic convulsive seizures would be obtained. This modification would greatly improve the use of the model to study TLE.

TLE is the most common type of focal epilepsy in adults (Thijs et al., 2019), with up to 30% of the patients achieving poor seizure control by currently available anti-seizure drugs (ASDs; (Kwan et al., 2011; Kapur et al., 2019). Moreover, several comorbidities challenge the quality of life of patients and their families (Keezer et al., 2016; Ravizza et al., 2017). As a result, it is important to facilitate research to identify new therapeutic targets, and to that end, animal models of epilepsy have played a key role (Holmes, 2007; Loscher, 2017; Pitkänen, 2017). Kainic acid (KA) has been used for decades in rats (Ben-Ari and Lagowska, 1978; Nadler et al., 1978; Schwarcz et al., 1978; Nadler, 1981; Ben-Ari and Cossart, 2000; Levesque and Avoli, 2013), where it can induce status epilepticus (SE), a pattern of neuronal loss similar to human TLE (mesial temporal sclerosis; MTS; (Houser, 1999; Scharfman, 2007; Blümcke et al., 2012; Thom, 2014), and spontaneous recurrent seizures (epilepsy), most of which are convulsive (Williams et al., 2007; Levesque and Avoli, 2013).

Although useful as a rat model of TLE, there often was mortality during or shortly after SE. Mortality was decreased by the use of anticonvulsants 1-2 hrs after SE (Scharfman et al., 2000; Scharfman et al., 2002) or by the use of repetitive low dosing (Meier and Dudek, 1996). However, there has been a major shift away from use of KA in rats to the use of KA in mice, a result of the powerful methods that were tailored for mice and can be used in epilepsy research.

Initial studies of mice often injected KA systemically. Unfortunately, initial studies in mice showed high mortality after systemic injection (Schauwecker and Steward, 1997). Use of anticonvulsants before KA led to reduced mortality (Iyengar et al., 2015) but the degree of hippocampal neuronal loss was sometimes weak even if electrographic SE lasted for several hours (Iyengar et al., 2015). In some studies, neuronal loss was greater (VonDran et al., 2014) but studies of long-term consequences showed that there were few spontaneous seizures (McKhann et al., 2003). Investigators tried to inject KA into the hippocampus instead and found pathology like MTS (Bouilleret et al., 1999). In addition, there was robust GC dispersion (GCD) which is also found in human TLE (Houser, 1992). The pathology following IHKA was useful because it simulated the cases of TLE where there is unilateral hippocampal sclerosis. In contrast, prior studies of systemic KA in rats produced severe pathology in both hippocampi. Another benefit of IHKA was low mortality (Bouilleret et al., 1999; Riban et al., 2002).

However, chronic convulsive seizures in the wks and months after IHKA in mice were not discussed in detail (Bouilleret et al., 1999; Riban et al., 2002) so the frequency and severity were not clear. EEG recordings from the site of IHKA injection showed that most frequent epileptiform abnormalities in mice were short-lasting non-convulsive episodes (Bouilleret et al., 1999; Maroso et al., 2011) which are not compelling seizures. Brief (3-7 sec) abnormalities were reported by others where they could be as frequent as 100 per hr (Kim et al., 2018; Sandau et al., 2019; Lai et al., 2020) which are also inconsistent with characteristics of seizures from an epileptic animal. Sometimes the EEG examples that were published suggested trains of spikes or epileptiform activity occurred in the wks after IHKA, but these are not clear seizures. Moreover, seizures were either rare, not shown, or the evidence that they occurred was not strong (e.g., (Kiasalari et al., 2016; Zhu et al., 2016; Runtz et al., 2018; Bielefeld et al., 2019; Li et al., 2020). One study that described methods for IHKA in mice concluded that IHKA did not lead to epilepsy (Bielefeld et al., 2017).

In addition to these problems, there have been other questions about the IHKA model in mice. For example, one lead is often used for the EEG and it is placed where KA was injected, without recording from other brain areas. This may lead to difficulty assessing how much the seizures spread beyond the hippocampus and the mistaken interpretation that seizures are focal. How many seizures were convulsive and how many were non-convulsive is not always clear. In addition, many studies occurred before the mandate at the National Institutes of Health (NIH) to study both sexes (Clayton and Collins, 2014), and this mandate is not in place outside the U.S. Therefore, most published data have used males.

To address these issues, we modified the IHKA procedure to produce frequent convulsive seizures while maintaining low mortality. We report the characteristics of these seizures that make them robust, such as multiple seizures per day or wk, severity in that most fit Racine scale 4-5, long duration (up to 100 sec), and postictal depression, a hallmark of convulsive seizures in TLE (So and Blume, 2010). We also report data for non-convulsive seizures and both sexes.

We also addressed other questions about IHKA in mice. For example, despite the report that post-IHKA epileptiform activity may change with time (Riban et al., 2002; Häussler et al., 2016), data about chronic seizures over time is limited, especially beyond 2 months post-IHKA (Henshall, 2017). Two studies mentioned in the text that convulsive seizures occurred many months after IHKA but data were not shown (Bouilleret et al., 1999; Bui et al., 2018). In contrast, quantified data from rats after systemic KA showed that spontaneous seizures increased over the first 6 months after IHKA injection from ∼9 per wk up to 50 per wk (Rattka et al., 2013). Therefore, we quantified seizures occurring 2-4 and 10-12 wks following IHKA injection. Remarkably, we show that seizure duration significantly increased. Although seizure frequency increased in approximately half of the mice, it declined in the other half, suggesting more investigators should monitor seizure duration to be sure a robust change with time is not overlooked.

Another question we addressed is the degree seizure types found in human TLE are also observed in the IHKA model. The seizures in humans have been characterized according to the pattern of the EEG during the initial phase of the seizure, called the seizure onset pattern. From a translational perspective, the identification of different seizure onset patterns is important because the type of seizure onset appears to correlate with the size of the seizure onset zone (Avoli et al., 2016) and may predict post-surgical outcomes (Zaher et al., 2020). The different seizure onset patterns are still being debated (Velasco et al., 2000; Perucca et al., 2014; Gnatkovsky et al., 2019; Saggio et al., 2020), but there appears to be a consensus that at least two types exist: low-voltage fast (LVF) or hypersychronous (HYP) seizures (Lévesque et al., 2012; Gnatkovsky et al., 2019). Although LVF and HYP seizures have been identified in IHKA-treated rats (Bragin et al., 2005), pilocarpine-treated rats (Behr et al., 2017), 4-aminopyridine or picrotoxin-treated rats (Salami et al., 2015) and slices of mice (Shiri et al., 2016), their presence in IHKA-treated mice has been unclear. This is an important gap because presently IHKA in mice is very common in studies of TLE. We show that almost all seizures in our IHKA-treated mice fit either the LVF or HYP classification.

In humans with TLE, high frequency oscillations (HFOs) are of great interest because they appear at seizure onset and may mark the best area for removal in surgical resection for intractable TLE (Zijlmans et al., 2012). Although observed in IHKA-treated rats (Bragin et al., 1999a), HFOs have not been described in epileptic mice to our knowledge. Furthermore, an important question is whether HFOs are confined to the focus, which is assumed to be the KA injection site in the IHKA model (Bouilleret et al., 1999; Riban et al., 2002). In addition, prior studies using one recording electrode at the injection site limited the assessment of seizure focus. We examined when HFOs occur, and where they occur.

In summary, we employed a slightly different approach to induce SE using IHKA and used continuous (24 hrs per day, 7 days per wk) wideband video-EEG (vEEG) monitoring from 4 recording electrodes at 2 different timepoints after IHKA. We asked several questions about our IHKA treated mice: 1) Were there frequent convulsive seizures? Did they generalize? Were non-convulsive seizures present? 2) Were seizures characterized by an onset pattern similar to LVF or HYP seizures? 3) Did seizures progress or remit? 4) Were there HFOs, and if so, where and when? Did HFOs mark a focus, and where was the focus? In addition, we examined hippocampal pathology to determine if it was similar to past reports of the IHKA model, i.e., MTS (Bouilleret et al., 1999). Finally, we made observations in both sexes to determine if there were sex differences.

## MATERIALS AND METHODS

### I. Animals, breeding, genotyping and animal care

The number of animals used for the present study is summarized in **Table 1**. All experimental procedures were performed in accordance with the NIH guidelines and approved by the Institutional Animal Care and Use Committee at the Nathan Kline Institute.

**Table 1.**
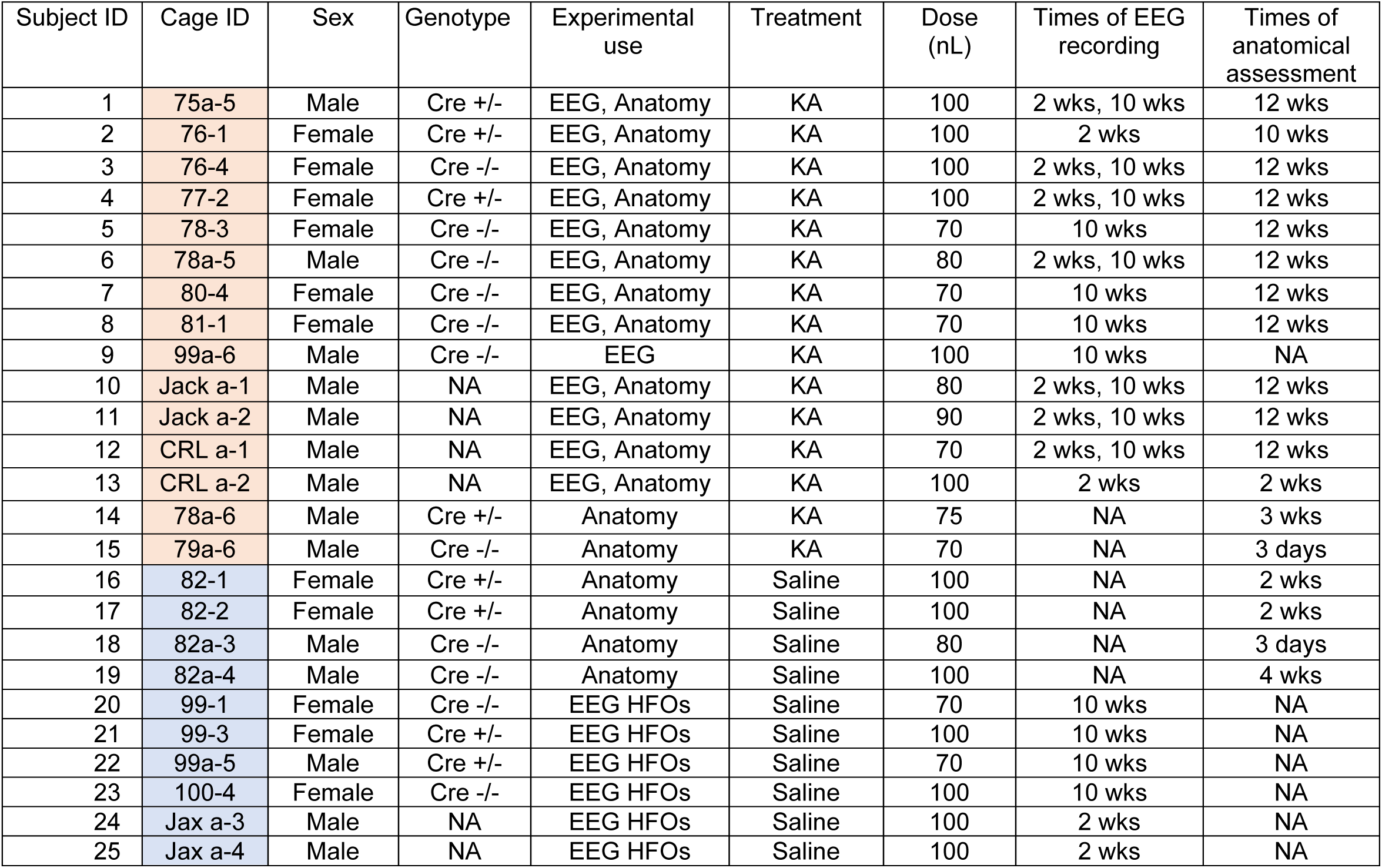
Animals included in the study. Animals included in the study. For each animal we note the assigned number, cage number (to show independent housing and what animals were from Jackson or Charles River Laboratories), sex, genotype, experimental use, treatment (KA, Saline), dose, times of 24 hr/day vEEG recording and anatomical assessment. The dose varied because pilot studies showed that doses between 70 and 100 nL all were successful in producing severe SE and there was no correlation between dose and the total number of chronic seizures (r=0.01, p=0.95). Therefore, doses were randomly assigned for this study. Orange: IHKA animals; Blue: Saline-injected controls. NA: Not applicable. Jack: Mice from Jackson Laboratories; CRL: Mice from Charles River Laboratories.

*Amigo2*-Cre transgenic mice were kindly provided by Dr. Steven Siegelbaum at Columbia University (Hitti and Siegelbaum, 2014). Hemizygous *Amigo2*-Cre males were bred to C57BL/6 females (Stock number 027, Charles River Laboratories). *Amigo2*-Cre+/- and *Amigo2*-Cre-/-adult males and females were used for experiments in anticipation of future closed-loop seizure intervention studies using this transgenic mouse line. Note that there was no effect of genotype on acute and chronic outcomes as described in more detail in **Table 2**. In addition, C57BL6 mice from the Jackson Laboratory (Stock number 000664) and Charles River Laboratories (Stock number 027) were used and no differences were found in acute and chronic epilepsy outcomes as explained in more detail in **Table 2**.

**Table 2.** Variables for induction of IHKA-SE and chronic epilepsy outcomes. For each animal we note the subject ID, sex, genotype, experimental use, treatment (KA, Saline), dose (nL of KA or Saline), total duration of anesthesia (defined as the time anesthesia started to the time anesthesia ended), latency to the first stage 5 seizure (defined as the time to a stage 5 convulsion after the end of anesthesia). We also note the total number of stage 5 seizures and chronic seizures (2-4 and 10-12 wk seizures were pooled). We also note whether animals were administered with 0.9% of Saline s.c. after IHKA (typically the day after IHKA i.e., 24 hrs post-IHKA). **IHKA dose:** There was no significant correlation between IHKA dose and the total number of stage 5 seizures during SE (r=0.19, p=0.5), suggesting the dose had little influence on the severity of SE. **Anesthesia:** There were no significant correlations between the total duration of anesthesia and the latency to the first stage 5 seizure (r=0.33, p=0.26), the total number of stage 5 seizures during SE (r=0.07, p=0.77), or the total number of chronic seizures (r=-0.47, p=0.1), suggesting that the duration of anesthesia had little influence on outcome after IHKA. **Latency to the first stage 5 and number of stage 5 seizures during SE:** Like the lack of correlation between anesthesia and latency to the first stage 5 seizure, there was no correlation between the latency to the first stage 5 and the total number of stage 5 seizures during SE (r=-0.23, p=0.39) or total number of chronic seizures (r=-0.38, p=0.19). These data suggest little effect of the latency of the first stage 5 seizures on the severity of SE or subsequent chronic seizures. The total number of stage 5 seizures during SE was not correlated with the total number of chronic seizures (r=-0.19, p=0.53) which is surprising because one might expect more severe SE to lead to more severe epilepsy. **Genotype:** We did not find any statistically significant differences between animals that were Cre -/- (n=7) or Cre +/- (n=4) in terms of the latency to the first stage 5 seizure (Mann-Whitney U-test, U = 8, p=0.31) or the total number of stage 5 seizures during SE (Mann-Whitney U-test, U = 10, p=0.50). Also, the total number of chronic seizures was not different between Cre -/- (n=6) vs. Cre+/- (n=3) animals (Mann-Whitney U-test, U = 8.5, p=0.96). These data suggest Cre-/- and Cre+/- mice were similar, regarding the measurements in this table. Also, the C57BL6 background strain was not different from Cre+/+ or Cre+/- mice in terms of the latency to the first stage 5 seizure (Cre+/+: Mann-Whitney U-test, U = 7, p=0.88, Cre+/-: Mann-Whitney U-test, U = 8, p=0.31) or the total number of stage 5 seizures during SE (Cre+/+: Mann-Whitney U-test, U = 2.5, p=0.14, Cre+/-: Mann-Whitney U-test, U = 10.5, p=0.59). Also, the mean number of chronic seizures in the background strain was not different from either Cre+/+ (Mann-Whitney U-test, U = 4, p=0.62) or Cre+/- (Mann-Whitney U-test, U = 8, p=0.47) mice. These data support the view that the use of either Cre+/+ or Cre+/- or the background strain led to no statistically significant effects in acute or chronic epilepsy outcomes examined in this study. **Administration of Saline:** Animals that received Saline s.c. the day after IHKA (n=3) because of transient body weight loss were similar to those that did not (n=10) in terms of total number of chronic seizures (Mann-Whitney U-test, U = 8, p=0.27). Orange: IHKA animals; Blue: Saline-injected controls. NA: Not applicable. Jack: Mice from Jackson Laboratories; CRL: Mice from Charles River Laboratories.

| Subject ID | Cage ID | Sex | Genotype | Experimental use | Treatment | Dose (nL) | Total duration of anesthesia (min) | Latency to Stg 5 (min) from IHKA injection | Total number of Stg5 during SE | Total number of chronic seizures | Administration of 0.9% Saline s.c. post-IHKA |
| --- | --- | --- | --- | --- | --- | --- | --- | --- | --- | --- | --- |
| 1 | 75a-5 | Male | Cre +/- | EEG, Anatomy | KA | 100 | 17 | 11 | 4 | 168 | No |
| 2 | 76-1 | Female | Cre +/- | EEG, Anatomy | KA | 100 | 17 | 28 | 8 | 8 | No |
| 3 | 76-4 | Female | Cre -/- | EEG, Anatomy | KA | 100 | 15 | 27 | 10 | 136 | Yes |
| 4 | 77-2 | Female | Cre +/- | EEG, Anatomy | KA | 100 | 15 | 42 | 6 | 59 | Yes |
| 5 | 78-3 | Female | Cre -/- | EEG, Anatomy | KA | 70 | 16 | 49 | 5 | 132 | No |
| 6 | 78a-5 | Male | Cre -/- | EEG, Anatomy | KA | 80 | 22 | 44 | 19 | 46 | Yes |
| 7 | 80-4 | Female | Cre -/- | EEG, Anatomy | KA | 70 | 19 | 66 | 4 | 8 | No |
| 8 | 81-1 | Female | Cre -/- | EEG, Anatomy | KA | 70 | 19 | 79 | 2 | 79 | No |
| 9 | 99a-6 | Male | Cre -/- | EEG | KA | 100 | 33 | 60 | 6 | 17 | No |
| 10 | Jack a-1 | Male | NA | EEG, Anatomy | KA | 80 | 21 | 17 | 5 | 44 | No |
| 11 | Jack a-2 | Male | NA | EEG, Anatomy | KA | 90 | 22 | 47 | 9 | 40 | No |
| 12 | CRL a-1 | Male | NA | EEG, Anatomy | KA | 70 | 18 | 30 | 8 | 54 | No |
| 13 | CRL a-2 | Male | NA | EEG, Anatomy | KA | 100 | 20 | 52 | 10 | 15 | No |
| 14 | 78a-6 | Male | Cre +/- | Anatomy | KA | 75 | 20 | 63 | 2 | NA | No |
| 15 | 79a-6 | Male | Cre -/- | Anatomy | KA | 70 | 17 | 39 | 8 | NA | No |
| 16 | 82-1 | Female | Cre +/- | Anatomy | Saline | 100 | 21 | NA | NA | NA | NA |
| 17 | 82-2 | Female | Cre +/- | Anatomy | Saline | 100 | 21 | NA | NA | NA | NA |
| 18 | 82a-3 | Male | Cre -/- | Anatomy | Saline | 80 | 21 | NA | NA | NA | NA |
| 19 | 82a-4 | Male | Cre -/- | Anatomy | Saline | 100 | 19 | NA | NA | NA | NA |
| 20 | 99-1 | Female | Cre -/- | EEG HFOs | Saline | 70 | 27 | NA | NA | NA | NA |
| 21 | 99-3 | Female | Cre +/- | EEG HFOs | Saline | 100 | 29 | NA | NA | NA | NA |
| 22 | 99a-5 | Male | Cre +/- | EEG HFOs | Saline | 70 | 28 | NA | NA | NA | NA |
| 23 | 100-4 | Female | Cre -/- | EEG HFOs | Saline | 100 | 31 | NA | NA | NA | NA |
| 24 | Jax a-3 | Male | NA | EEG HFOs | Saline | 100 | 21 | NA | NA | NA | NA |
| 25 | Jax a-4 | Male | NA | EEG HFOs | Saline | 100 | 22 | NA | NA | NA | NA |

Prior to IHKA injection, mice were housed 2-4 per cage in standard laboratory cages and after IHKA injection they were housed 1 per cage. Animals were handled daily by the experimenter when they were single housed to reduce behavioral stress related to the lack of social housing (Bernard, 2019; Manouze et al., 2019). Cages were filled with corn cob bedding and there was a 12 hr light:dark cycle (7:00 a.m. lights on, 7:00 p.m. lights off). Red plastic mini igloos (W.F. Fisher) were placed at the base of the cage to provide a location that was partially hidden for the mice. This was done to reduce behavioral stress. Food (Purina 5001, W.F. Fisher) and water was provided ad libitum.

All breeding pairs were fed Purina 5008 rodent chow (W.F. Fisher) and provided with one 2’’x 2’’ nestlet (W.F. Fisher). Mice were weaned on postnatal day 23-30, and after that time the chow was Purina 5001 (W.F. Fisher).

Genotyping was done using tail samples collected at approximately 23-30 days of age. Genotyping was performed by the Mouse Genotyping Core Laboratory at New York University Langone Medical Center.

Although this study used *Amigo2*-Cre mice we do not think the presence of Cre recombinase in cells expressing *Amigo2* influenced the results. One reason is that our results from *Amigo2*-Cre+/- and *Amigo2*-Cre-/- mice were similar in several acute (during SE) and chronic (during chronic seizures) outcome measures as described in more detail in **Table 2**. In addition, *Amigo2* has been localized only to CA2 pyramidal cells and hilar neurons (Hitti and Siegelbaum, 2014), which is a small fraction of neurons in the brain. However, it is notable that in studies of others, *DLX*-Cre+/-mice did appear to have more seizures than *DLX*-Cre-/- mice despite the absence of viral injection to experimentally manipulate cells expressing *DLX* (Kim et al., 2013). Other laboratories have used Cre+/- lines without viral expression in their research, however, and it has not been shown that hemizygous Cre on its own has major effects on endpoints we investigated, e.g., seizures after SE.

### II. Kainic acid (KA) injection

#### A. KA preparation

KA monohydrate (#0222, Tocris Bioscience) was dissolved in 0.9% sterile Saline and pH was adjusted to 7.4 with 20-30 µl 1 N NaOH (#SS266-1, Fisher Scientific) according to Tocris Bioscience’s guidelines. The final concentration of the stock solution was 20 mM and it is similar to the concertation used by other investigators (Bui et al., 2018; Zeidler et al., 2018; Li et al., 2020). The stock solution was then sonicated for 1 hr to ensure good solubility as indicated in the manufacturer’s recommendations. After sonication, the solution was aliquoted in 0.5 ml portions and kept at -80°C for a maximum of 1 month. On the day of IHKA injection, an aliquot was allowed to come to room temperature, and the solution was sonicated for 30 min before the start of the surgical procedure. The aliquot was discarded after use and a new aliquot was used on the day of every experiment. Using this approach, we did not notice any precipitation at any step of KA preparation.

#### B. Stereotaxic injection of KA

At approximately 8 wks of age, mice were injected with KA and then implanted with electrodes 10 days or 9 wks later (see methods for KA injection in the next paragraph and implantation in section IV), which was 4-7 days before starting vEEG monitoring. Although implantation at 10 days after IHKA may have altered epileptogenesis, animals exhibited spontaneous seizures in their home cage by 10 days after IHKA, so epilepsy had already developed. Furthermore, in our experience, implantation reduces seizures rather than increasing them (Jain et al., 2019), so it is likely that our results underestimate (rather than overestimate) seizures. This is important because our goal was to show robust seizures rather than few, equivocal seizures. Note that implantation prior to IHKA injection would have potentially allowed detailed recordings of SE, but that was not the major focus of this study. Moreover, prior implantation can reduce seizures as mentioned above, so it was avoided.

To begin the procedure, the 8 wk-old mice were brought to the laboratory for acclimation to the location where KA would be injected. Acclimation typically included two 5 min-long sessions per day for the 2 days before IHKA injection. In each session, the investigator who would be injecting KA conducted the acclimation. For acclimation, the mouse was removed from the cage using gloves and the mouse was allowed to walk on the part of the lab coat covering the lower forearm. One M&M (chocolate-coated peanut) was used as a food reward.

KA was injected between 8 a.m.-1 p.m. Mice were anesthetized with 3% isoflurane (Aerrane, Henry Schein) for 2 min in a rectangular transparent plexiglass box (induction chamber) and then transferred to a stereotaxic apparatus (Model #502063, World Precision Instruments). Anesthesia was then lowered to 1-2%. Mice were frequently monitored to confirm there was adequate respiration and there was no reflex in response to a toe pinch. Mice were placed on top of a homeothermic blanket (Model #50-7220F, Harvard Apparatus) and body temperature was maintained at 37°C by feedback from a lubricated probe inserted into the rectum. Eye ointment was applied to prevent dehydration (Artificial tears, Pivetal). The scalp was shaved and swabbed with Betadine (Purdue Products) using sterile cotton-tipped applicators (Puritan) followed by 70% ethanol. A midline incision exposing the skull surface was made with a sterile scalpel and the skull was cleaned with sterile Saline.

One burr hole was drilled using a drill bit (Model #514552, 60 mm, Stoelting) mounted to a surgical drill (Model C300, Grobert) above the left hippocampus. Stereotaxic coordinates for the burr hole were (-2 mm anterior-posterior (A-P) to Bregma, - 1.25 mm medio-lateral (M-L)). Care was taken to leave the dura mater intact by regularly monitoring it with a stereoscope during drilling (Stemi SV6, Zeiss). The drilling area was regularly hydrated with 0.9% sterile Saline solution and drilling was done in steps to avoid depressing the underlying tissue during drilling. This approach was followed because we measured the dorso-ventral (D-V) zero point from dural surface, so we wanted the measurement of the dural surface to be as accurate as possible.

Next, a 0.5 ml Hamilton syringe (Model 7001, Hamilton) was lowered from brain surface 1.6 mm into the left dorsal hippocampus (-1.6 mm D-V) and 70-100 nL of 20 mM KA dissolved in 0.9% sterile Saline was manually injected over approximately 5 min. A range of IHKA volumes were used instead of a fixed volume because in pilot studies all doses were successful in triggering robust convulsive SE with minimal (approximately 20%) acute (during SE) mortality (see the legend for **Table 2** for a detailed explanation). The speed was controlled by depressing the syringe 1 unit (10 nL) every 15 sec. The needle remained in place for an additional 3 min and was then slowly removed to prevent backflow of the injected solution. The incision was quickly closed using tissue adhesive (Vetbond, 3M). Mice were transferred to a clean cage on an autoclaved paper towel (without bedding) and placed over a 37°C heating pad. After approximately 3-4 hrs of monitoring the animal, bedding was used instead of the towel.

### III. Behavioral monitoring of IHKA-induced status epilepticus (SE): convulsive and non-convulsive seizures

Behavior during IHKA-induced SE was visually monitored and seizure severity during SE was scored using stages 1-5 of the Racine scale (Racine, 1972) and stages 6-7 using the Pinel and Ronver scale (Pinel and Rovner, 1978) because all stages were observed in our experiments. Using the scale with stages 1-7, stages 1-2 were considered **non-convulsive** and stages 3 and higher were considered **convulsive.** Stage 1 was accompanied by intense mastication and facial movements such as blinking, repetitive ear movements, a sudden change in the behavior to a frozen stance. Stage 2 was head nodding. Stage 3 was unilateral forelimb clonus, stage 4 was bilateral forelimb clonus with rearing, and stage 5 was a stage 4 seizure with loss of postural tone. Stage 6 seizures were accompanied by jumping or repetitive falling and stage 7 included vigorous jumping and running around the cage. Notably, all IHKA-injected animals experienced vigorous convulsive SE meaning that there were several stage 5 seizures. After the first stage 5 seizure, the cage was removed from the heating pad to prevent hyperthermia. The body temperature was regularly (approximately every 15 min) monitored with an infrared thermometer (Model #800048, Sper Scientific) to ensure that body temperature was within the physiological range (36-37°C).

Mice were visually monitored until normal behavior (defined as exploration and/or grooming) resumed. After 3-4 hrs of observation following IHKA injection, mice resumed normal behavior, and they were returned to their home cages with normal bedding and no paper towel. The cages were kept overnight in the same location. After 1 day, mice were moved close to the EEG equipment so that they would acclimate to that area.

Starting with the day after IHKA injection, animals were handled for approximately 5 min daily as described above and their body weight was measured for the next 7 days once daily. In 3 IHKA-injected mice (1 male, 2 female), body weight declined transiently by 10-18% so they were administered 3 ml of warm (36-37°C) sterile Saline solution s.c. which did not affect chronic outcome measures as described in **Table 2**. The solution was administered once and within 24 hrs post-IHKA. These and other aspects of the IHKA injection procedure and subsequent 7 days are listed in **Table 2**.

### IV. EEG Recording

#### A. Surgical implantation of EEG electrodes

Two wks after IHKA injection, animals were implanted with 6 subdural electrodes (4 recording, 1 reference, 1 ground; see below for details). As mentioned above, this time was chosen because animals were already exhibiting convulsive seizures in their homecage, so the period of epileptogenesis appeared to be over.

Mice were anesthetized with 3% isoflurane (Aerrane, Henry Schein) for 2 min and then transferred to the stereotaxic apparatus (Model #502063, World Precision Instruments). Anesthesia was then lowered to 1-2% and monitored the same way as described above for IHKA injection. Note that vendor information is already specified above for many of the items used below and where not the vendor information is provided. Mice were placed on top of a homeothermic blanket, eye ointment was applied, and the scalp was shaved and cleaned following the same procedures described above. A midline incision was made with a sterile scalpel and the skull was cleaned with sterile Saline. Two burr holes were drilled over the left and right hippocampus (-2.5 mm A-P, ± 1.75 mm M-L), slightly posterior to the septotemporal level where KA was injected. Two additional burr holes were drilled anterior to the IHKA injection site (-0.5 mm A-P, ± 1.5 mm M-L) to serve as the cortical recording leads. Subdural screw electrodes (0.10’’ length stainless steel) were placed in the burr holes and secured using dental cement (Jet Set-4 Denture Repair, Lang Dental). Two additional burr holes were drilled above the cerebellar region to serve as reference (-5.7 mm A-P, +1.25 mm M-L) and ground (-5.7 mm A-P, -1.25 mm M-L). The reference was placed contralateral to the IHKA injection site and the ground was ipsilateral. The subdural screw electrodes were attached to an 8-pin connector (#ED83100-ND, Digikey) which was secured to the skull with dental cement.

#### B. Continuous wide-band video-EEG monitoring

One-day after EEG surgery and until the vEEG recording started, mice were housed in the room where the vEEG equipment was located so that they could acclimate to the recording environment. For recording, mice were placed into a 21 cm x 19 cm square transparent plexiglass cage which had access to food and water and had no cage lid.

Food was placed on the bottom of the cage and a water bottle was attached to one of the walls of the cage. A pre-amplifier (Pinnacle Technologies) was connected to the 8-pin connector and then to a commutator (Pinnacle Technologies) which allowed free movement of the mouse in the entire cage. EEG signals were amplified 10 times and recorded at 2 kHz sampling rate using a bandpass filter (0.5-500 Hz) and Sirenia Acquisition System (Pinnacle Technologies). High frame rate video (30 fs) was recorded simultaneously using an infrared LED camera (Model #AP-DCS100W, Apex CCTV).

#### C. Quantification of video-EEG

##### 1. Seizure analyses

Seizures were detected from vEEG by a blinded investigator (CPL) using replay of the EEG and video records. The left hippocampal lead was chosen for most measurements because it was the site of the IHKA injection. However, seizures were typically present in all leads. Therefore, they demonstrated robust synchronization and generalization.

**The electrographic component of a seizure**, whether convulsive or non-convulsive, was defined by a sudden change in amplitude >2x of the standard deviation (SD) of the baseline mean. The threshold, 2x the SD of the baseline mean, was chosen because it was adequate to differentiate seizures from normal EEG. The baseline mean was calculated from the 30 sec prior to the event that was being considered to be a possible seizure. During this 30 sec baseline, care was taken to choose a time when the behavioral state was not interrupted by large artifacts or abrupt behavioral state changes. **An electrographic seizure** was also defined by high frequency rhythmic activity (>5 Hz) which consisted of an abnormal pattern (large amplitude spikes and clusters of spikes lasting for at least 10 sec). Ten seconds was chosen because seizures in TLE typically last at least 10 sec and often are 20-60 sec (Balish et al., 1991). In addition, most seizures recorded in a standard Epilepsy Monitoring Unit typically last more than 10 sec (Jenssen et al., 2006). We also have found that the majority of seizures in epileptic mice after pilocarpine-induced SE in our laboratory were at least 10 sec and usually 20-60 sec (Iyengar et al., 2015; Botterill et al., 2019; Jain et al., 2019). **Seizure onset** was defined as the time when the baseline of the left hippocampal lead exceeded 2x the SD of the baseline mean. The **end of a seizure** (seizure termination) was defined as the time when high amplitude activity declined to <2x the SD of the baseline mean. **Seizure duration** was calculated by subtracting the time of seizure termination (the end of the seizure as defined above) from the time of seizure onset. **Seizure burden** was quantified as the number of days that animals sustained chronic seizures (either non-convulsive or convulsive) divided by the total number of days that continuous vEEG was performed at 2-4 and 10-12 wks (i.e., 14 days). For each seizure, the light cycle was noted as lights on (7:00 a.m.) or off (7:00 p.m.) and the time of the day with a.m. and p.m. The behavioral state when the seizure began was determined as either wakefulness or sleep based on video and the EEG for a period of 30 sec before seizure onset. The **awake** state included awake exploration and awake immobility. **Awake exploration** was defined by walking or other movement around the cage and the predominance of theta rhythm (4-12 Hz in the hippocampal leads) with eyes open (when the mouse was facing the camera). **Awake immobility** was not accompanied by exploration, but eyes were open. **Sleep** was defined either as non-rapid eye movement (NREM) or rapid eye movement (REM) sleep. **NREM** was dominated by slow wave activity in the delta (<5 Hz) frequency range and eyes were closed with no movement of the body. **REM** sleep was defined by theta rhythm (4-12 Hz) and eyes closed with no movement of the body. For defining sleep stages, we relied on leads contralateral to the IHKA injection site because of previous reports about loss of theta rhythm predominantly adjacent to the IHKA injection site (Riban et al., 2002; Arabadzisz et al., 2005).

For quantification, the percent of seizures in each category (on or off, a.m. or p.m.; awake or asleep) was calculated by dividing the number of seizures in a category by the total number of seizures. The total was either the total number per animal or the total number of all seizures in all animals.

Seizure onset patterns were defined LVF, HYP (as described above) or ‘unclear’ which was a seizure that had an onset pattern that was not possible to classify as LVF or HYP. LVF seizures started with a sentinel spike (amplitude >2x SD above the baseline mean) followed by brief suppression of the background EEG whereas HYP seizures began with high amplitude (>2x SD above the baseline mean) spiking at a frequency that was much greater than interictal spikes. These definitions are similar to those described in other animal models of epilepsy (Bragin et al., 2005; Behr et al., 2017) and human patients (Velasco et al., 2000).

##### 2. High frequency oscillations (HFOs)

**HFOs** were defined as oscillations >250 Hz consistent with past studies in humans (Staba et al., 2004) and animals (Bragin et al., 1999a). To sample HFOs, NREM sleep was selected because prior studies suggested that HFOs occur mainly during slow waves and sharp waves occurring in NREM sleep (Staba et al., 2004; Bagshaw et al., 2009). For each animal, an interictal period was chosen, and 10 min was sampled at least 1 hr after the last seizure and at least 1 hr before the next seizure. All 4 channels of wideband interictal EEG were visually inspected for the presence of HFOs and a 10 min NREM sleep epoch was selected. This was possible because high frequency activity was visually discernable from the ongoing background EEG activity. We then applied an automated approach to detect HFOs using the RippleLab application written in MATLAB (Navarrete et al., 2016) and the algorithm developed by Staba and colleagues (Staba et al., 2002). Automated detection was followed by visual inspection of putative HFOs and inclusion/exclusion criteria were used to either include or exclude the putative events as HFOs. These criteria are described below. In brief, each channel was band-pass filtered between 250-500 Hz using a finite impulse response (FIR) digital filter and the root mean square (RMS) amplitude of the filtered signal was quantified. Successive RMS amplitudes >5x SDs above the mean amplitude of the RMS signal calculated for the entire 10 min epoch (during NREM) that lasted >6 msec in duration were defined as putative HFOs. Putative HFOs that did not have at least 4 peaks in the rectified band-passed signal >3x SDs greater than the mean of the baseline signal were excluded. We then reviewed each putative HFO to exclude artifactual events associated with movement or other sources of noise (Bénar et al., 2010; Menendez de la Prida et al., 2015; Amiri et al., 2016). To better appreciate HFO power in time, we applied time-frequency analyses to visualize HFO power. To that end, we used the time-frequency function which is part of RippleLab (Navarrete et al., 2016) and applied a 64-point window to analyze HFO power in the 250-500 Hz frequency domain.

Whether HFOs were interictal or ictal was visually determined by inspecting all 4 channels. **Interictal HFOs** were determined from the EEG at least 1 hr before or 1 hr after a seizure. **Ictal HFOs** were visually defined as HFOs that occurred at seizure onset and during seizures by inspecting all 4 channels for the presence of HFOs >250 Hz.

### V. Anatomy

#### A. Perfusion-fixation and sectioning

Mice were deeply anesthetized with isoflurane and they were injected with an overdose of urethane (2.5 g/kg, i.p.; Sigma Aldrich). During surgical anesthesia, defined as the time when mice did not respond to a toe pinch with a withdrawal reflex, the abdominal cavity was opened with a scalpel and a 25 gauge butterfly-style needle (Model #J454D, Jorvet) was inserted into the heart. When the needle was clamped in place with a hemostat, mice were transcardially perfused with 10 ml of room temperature (25°C) Saline, followed by 30 ml of cold 4% paraformaldehyde (#1920, Electron Microscopy Systems) in 0.1 M phosphate buffer (PB; pH=7.4) using a peristaltic pump at a rate of 10-12 ml/min (Minipuls2, Gilson). The brain was quickly removed and stored overnight in 4% paraformaldehyde in 0.1 M PB. An incision was made laterally on the right side of the brain to mark the side that was contralateral to the IHKA injection. On the next day, the brains were sectioned (50 µm thickness) in the coronal plane using a vibratome (Vibratome 3000, Ted Pella) in Tris buffer. Sections were stored in 24 or 48 well tissue culture plates containing cryoprotectant solution (25% glycerol, 30% ethylene glycol, 45% 0.1 M PB, pH 6.7) at 4°C until use.

#### B. Nissl staining

For Nissl staining, sections were mounted on 3% gelatin-coated slides and left to dry overnight at room temperature. Then slides were dehydrated in increasing concentrations of ethanol (70%, 95%, 100%, 100%) for 2.5 min each, cleared in Xylene (Sigma Aldrich) for 10 min, and then dehydrated again (70%, 95%, 100%, 100%) followed by hydration in double-distilled (dd) H_2_0 for 30 sec. Then sections were stained with 0.25% cresyl violet (Sigma Aldrich) dissolved in ddH_2_0 for 1.5 min followed by 30 sec in 4% acetic acid solution dissolved in ddH_2_0. Then sections were dehydrated (70%, 95%, 100%, 100%), cleared in Xylene for 4 min, and cover-slipped with Permount (Electron Microscopy Systems).

#### C. Quantification

To estimate neuronal loss and granule cell dispersion in dorsal hippocampus, we first chose one coronal section per animal that was selected slightly anterior to both the IHKA or Saline injection site. This site also allowed us to assess tissue integrity near the injection track and subdural electrode near the injection site. This dorsal section corresponded to approximately -1.94 to -2.06 mm posterior to Bregma in a common mouse stereotaxic atlas (Figures 47-48; (Franklin and Paxinos, 1997). We chose this level to conduct initial quantification because previous reports suggesting that neuronal loss is maximal adjacent to the IHKA injection site (Bouilleret et al., 1999; Riban et al., 2002). In both IHKA- and Saline-injected mice, good tissue integrity was observed near the injection site, and the injection track was actually hard to see. This absence of damage by the injection needle rendered anatomical assessments possible at this tissue level. The absence of a visible injection track was probably achieved because we used a syringe and not a permanently implanted (guide) cannula to inject KA or Saline.

The absence of damage from the EEG recording electrode was because we used subdural screw electrodes instead of depth electrodes for our vEEG monitoring. All these approaches significantly minimized tissue damage near the injection site, providing an opportunity to assess the injection area with limited technical confounds.

After this initial assessment, we added hippocampal sections that were more posterior to the IHKA injection site (-2.5 mm to Bregma). That was done to determine if neuropathology is evident beyond the IHKA injection site. Importantly we found the damage was present posteriorly, so it extended beyond the IHKA injection site, which is not always reported.

Nissl-stained sections were imaged using a microscope (Model BX51, Olympus of America) and digital camera (Infinity3-6URC) at 2752x2192 pixel resolution. Photographs were taken with Infinity capture software (version 6.5.6). After images were imported to ImageJ software (NIH), we measured the pyramidal cell layer length (PCL length), granule cell layer (GCL) area, and GCL width or thickness. Details are shown in **Supplemental Figure 1**. For PCL length, a line was drawn using the freehand line tool from the border of the PCL in CA3c with the DG hilus to the CA1 border with the subiculum. The CA3c cell layer at the border with the hilus was defined as the point where pyramidal cells became sufficiently close so that there was <1 pyramidal cell body width apart from adjacent PCs. The CA1 cell layer at the border with the subiculum was defined in an analogous manner because the cell layer broadens significantly as the subiculum begins. For GCL area measurements, we traced the borders of the GCL using the freehand tool. The GCL was distinguished from the hilus as the point where GC somata had <1 cell width distance between them. For GCL thickness, we used the straight-line tool to measure the thickest part of the layer. The line to measure thickness was made perpendicular to the GCL, starting at the border with the hilus and ending at the border with the inner molecular layer (IML). Two measurements were made of the widest part of the upper and the lower blade, and then we averaged the 2 measurements.

### VI. Statistics

Data are presented as mean ± standard error of the mean (SEM). Statistical significance was set at p < 0.05 and is denoted by asterisks on all graphs. All statistical analyses were performed using Prism (version 8.4.2, Graphpad). Comparisons of parametric data of two groups were done using unpaired or paired two-tailed Student’s t-test. To determine if data fit a normal distribution the D’Agostino-Pearson test was used in Graphpad. To determine if the variance of groups was homogeneous, the Brown-Forsythe test was used in Graphpad. When data did not fit a normal distribution or transformation was unable to resolve differences in variance, nonparametric statistics were used. The non-parametric tests were the Mann-Whitney U-test to compare two groups and the Kruskal-Wallis ANOVA for multiple groups. For correlation analyses we used linear regression and the Pearson correlation coefficient (r) in Graphpad.

### VII. Experimental design and data collection

The present study used *Amigo2*-Cre+/- or *Amigo2*-Cre-/- mice for IHKA and Saline injections. In addition, four C57BL6 mice from the Jackson Laboratory (n=2; Stock number 000664) or Charles River Laboratories (n=2; Stock number 027) were used as controls for the background strain. Animals were first acclimated and then KA or Saline was injected in the hippocampus. A cohort of 8 IHKA-injected animals was recorded at 2-4 wks post-IHKA and the same mice were recorded again at 10-12 wks with continuous wideband vEEG monitoring. In addition, 4 IHKA-injected animals were recorded only at 10-12 wks. Six animals were injected with Saline and EEG was recorded as controls for HFOs and to confirm a lack of seizures.

For anatomy, animals were perfusion-fixed after vEEG, 12 wks post-IHKA (n=10). Earlier times were also checked to confirm a similar degree of PCL damage (3 days, n=1; 3 wks, n=1), consistent with the idea that most pyramidal cell loss occurs shortly after SE (Sutula and Pitkanen, 2002). Saline-treated mice were examined also (3 days, n=1; 2-4 wks, n=3) and did not show neuronal loss, as described in more detail in the Results. A list of all animals included in the study is presented in **Table 1**. Only one animal died in this study, and that was unrelated to the study.

### VIII. Data collection

Although, EEG analyses and quantification of Nissl-stained sections were done by an investigator (CPL) who was blinded to the experimental group and sex, data collection was unblinded because animals were more active after IHKA injection compared to Saline-injected controls.

Data collection was done using Case Report Forms (CRFs), where each CRF was specific for a procedure such as IHKA injection. Common Data Elements (CDEs) were compiled as previously described (Harte-Hargrove et al., 2018; Scharfman et al., 2018). We provide all CDEs we used in each procedure-specific CRF as a supplemental .xls file.

## RESULTS

### I. IHKA leads to epilepsy with frequent spontaneous convulsive seizures

To determine whether convulsive seizures occurred in the mice that were injected with KA (IHKA-treated mice), mice were monitored by continuous vEEG from 2-4 and 10-12 wks after IHKA-induced convulsive SE (**Figure 1A**). At both the early (2-4 wks) and late (10-12 wks) times, convulsive seizures occurred spontaneously and were typically frequent (multiple seizures per day) and severe (stage 4-5). A representative example of a convulsive seizure is shown in **Figure 1B1** and is expanded in **Figure 1B2**. During the part of the EEG that appeared to be a seizure, convulsions typically developed in the middle or later parts of the EEG seizure activity. Thus, the EEG showed evidence of a robust seizure first, and then a convulsion developed before the EEG manifestations of the seizure ended. At the same time as loss of postural tone, or after the convulsive behavior ended, the EEG manifestations of the seizure ended.

**Figure 1.**
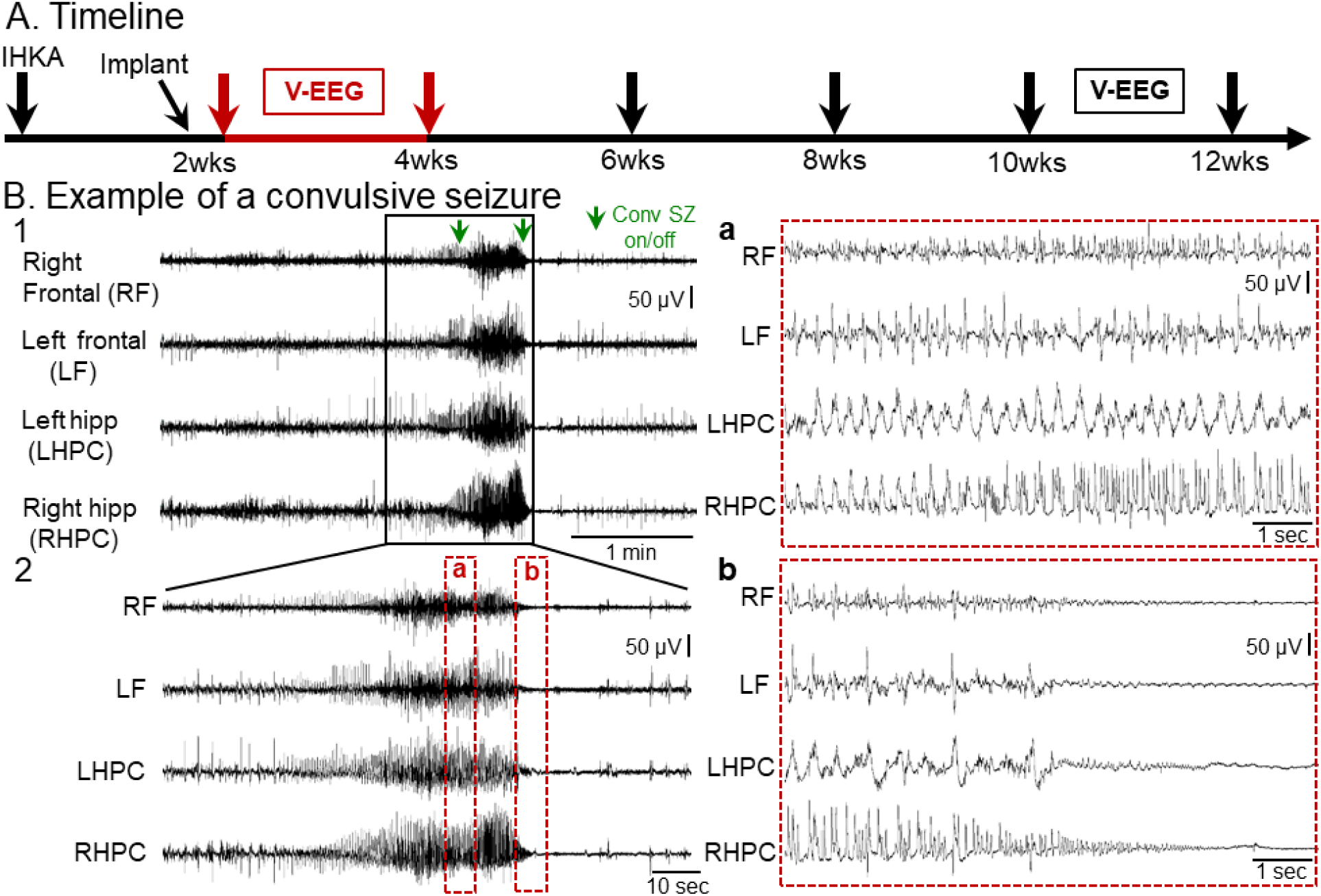
Example of a spontaneous convulsive seizure recorded 2-4 wks after IHKA. **(A)** Experimental timeline of the study. Animals were injected with KA in the left dorsal hippocampus and 10 days later they were implanted with 4 subdural screw electrodes. They were then vEEG monitored 2-4 wks post-IHKA continuously (red arrows, with data from this time shown in B) and then recorded again at 10-12 wks post-IHKA. **(B)** Representative example of a chronic convulsive seizure recorded during the 2-4 wk session after IHKA. Four leads were used to record right and left frontal cortices and left and right hippocampal regions. A 5 min-long EEG trace with seizure activity in all 4 leads is shown in B1. The start and the end of the convulsive seizure is indicated by green arrows (Conv SZ on/off). The same seizure is shown in a 2 min-long time window in B2 and further expanded (a, b) in 10 sec epochs to better show the EEG complexity. Inset a of the seizure in B2 shows complex and often rhythmic activity with fast and slow components. Inset b of the seizure in B2 shows prolonged suppression of the background EEG in all 4 leads after the termination of the electrographic manifestations of the seizure.

Next, we quantified chronic spontaneous convulsive seizures during the 2-4wks post-IHKA (**Figure 2A**) and results are shown in **Figure 2B-H**. The total number of convulsive seizures during the 2 wks is shown in **Figure 2B**, highlighting that every animal sustained multiple convulsive seizures. A breakdown of the number of seizures per day showed some variability between animals, although all animals had multiple convulsive seizures per day for many days in a row (**Figure 2C**). Note that one of the females (76-1) had fewer seizures than other mice (**Figure 2B-D**), but that mouse did have 5 convulsive seizures in 2 wks which is still a clear demonstration of chronic epilepsy.

**Figure 2.**
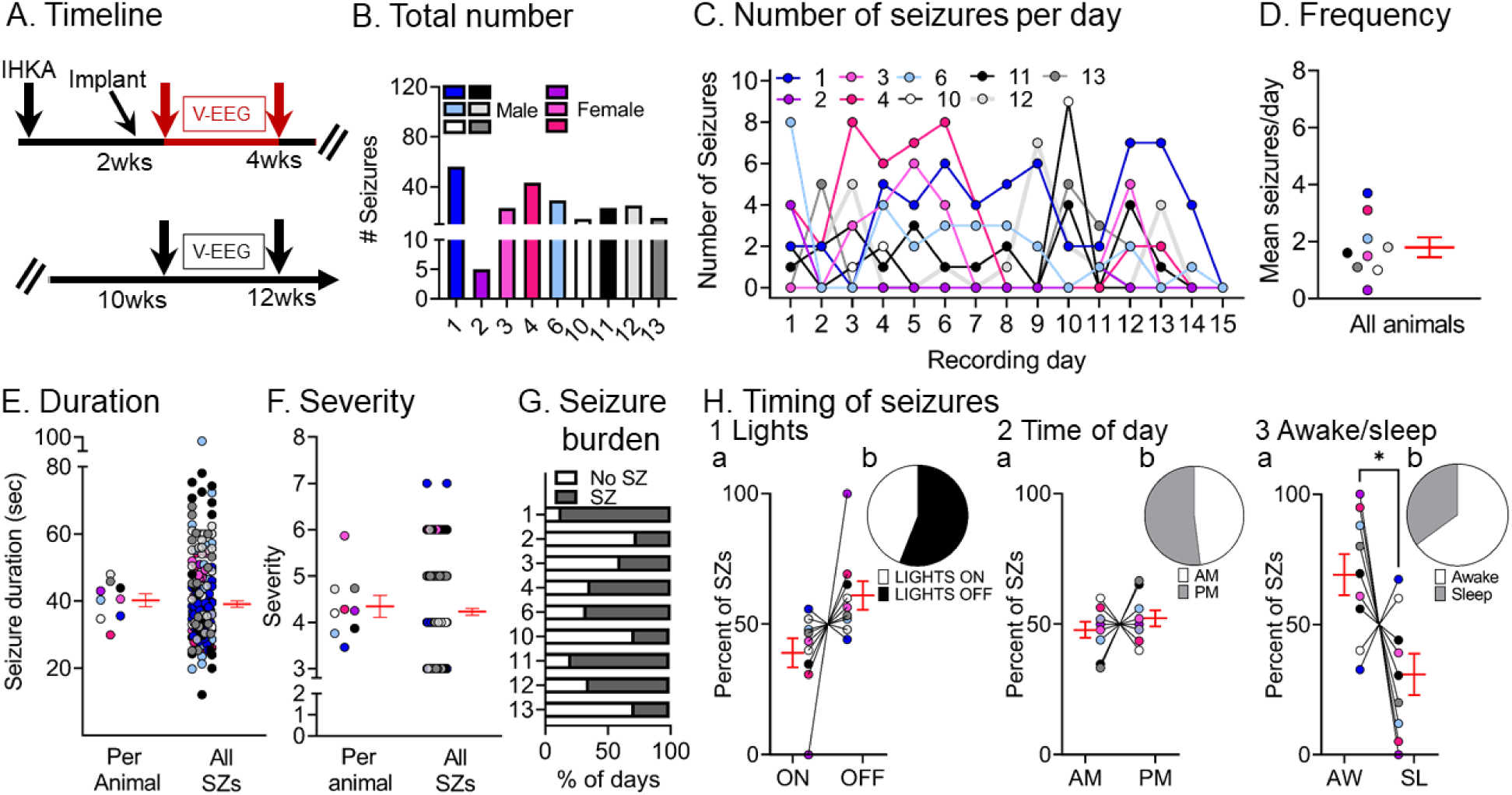
Quantification of chronic spontaneous convulsive seizures 2-4 wks post-IHKA. **(A)** Experimental timeline of the study. The 2-4 wk data were used in this figure (red arrows). For this timepoint, there were 9 mice (males, blue, gray, or white; females, pink; total # seizures=233). **(B)** The total number of chronic convulsive seizures recorded 2-4 wks post-IHKA is shown per animal. Note that animals showed frequent convulsive seizures although there was variability. One of the female mice, 76-1, had 5 seizures in 2 wks which is fewer than other animals, but nevertheless is a demonstration of chronic epilepsy. **(C)** The number of convulsive seizures is plotted for each day of the recording period. Each animal is a different line, has a different color designation, and was assigned a different number. The color coding is the same in subsequent figures to make it possible to compare each animal across figures. **(D)** Convulsive seizure frequency was calculated as the mean number of convulsive seizures per day for each animal. Data are presented in D-F and H as individual values and as mean ± SEM (red). **(E)** Convulsive seizure duration was calculated as the mean per animal (left) or the mean of all convulsive seizure durations (right; n=211 seizures). **(F)** Convulsive seizure severity was calculated as the mean per animal (left) or the mean of all convulsive seizures (right; n=211 seizures). **(G)** Convulsive seizure burden was defined as the percent of days spent with (SZ) or without (No SZ) seizures. **(H)** The percent of convulsive seizures is shown, either occurring during the light period or dark period of the light:dark cycle (Lights ON or OFF; H1), a.m. or p.m. (AM, PM; H2), and in awake (AW) or sleep (SL) state (H3). The percentages were calculated as the mean per animal (H1a, H2a, H3a) or the mean of all seizures (H1b, H2b, H3b). There were no significant differences (paired t-tests; all p>0.05) for the light:dark cycle or a.m. vs. p.m. However, a significantly higher percentage of seizures occurred during awake vs. sleep stage (paired t-test, *t*_crit_ = 2.416, p=0.04).

The mean convulsive seizure frequency was 1.8±0.35 seizures per day (range 0.3-3.7, n=9 mice; **Figure 2D**). Convulsive seizures lasted for 40.22±1.94 sec (range 29.9-47.9) when mean duration was calculated per animal, and when all seizures were pooled, the mean seizure duration was 39.1±0.8 sec (range 12.08-94.5; **Figure 2E**).

Analyses of simultaneous video records revealed that convulsive seizures fit the modified Racine scale with a mean severity score of 4.3±0.2 (range 3-6) when the mean was calculated per animal, and 4.2±0.07 (range 3-7) when all seizures were pooled (**Figure 2F**). Next, we quantified the number of days that animals sustained convulsive seizures in 2-4 wks because this metric gives us an estimate of seizure burden (**Figure 2G**). The mean percent of days spent with seizures (53.7±7.7%, range 27-87%) was not different from the days spent without seizures (45.9±7.7%, range 13-73%; paired t-test, *t*_crit_ = 0.503, p=0.62). In order to account for any potential circadian effects on seizures, we calculated the percent of seizures occurring during the 12 hr-long period when lights were on and the 12 hr-long period when lights were off (**Figure 2H1a, 2H1b**), and no significant differences were found (paired t-test, *t*_crit_ = 1.973, p=0.08). We also did not find any differences when we examined the a.m. or p.m. (12:00 a.m.-12:00 p.m. vs. 12:00 p.m. to 12:00 a.m.; paired t-test, *t*_crit_ = 0.726, p=0.48; **Figure 2H2a, 2H2b**). Finally, we examined the times when mice were awake and times they were asleep (awake and sleep are defined in the Methods) and we found a higher percentage of seizures occurring at the awake state (paired t-test, *t*_crit_ = 2.416, p=0.04; **Figure 2H3a, 2H3b**).

In summary, convulsive seizures were robust both qualitatively and quantitively in IHKA-treated mice. Seizures were typically severe and observed with all recording electrodes, indicating that they were generalized. They also showed additional characteristics that have been discussed in human TLE, including HFOs, which are discussed further below.

### II. Chronic convulsive seizure frequency varies over time

Next, we continued to ask whether our IHKA model was robust. To that end, we determined whether convulsive seizures persisted at 10-12 wks, 2.5-3.0 months after IHKA.

Seven out of 9 animals recorded at 2-4 wks were recorded again at 10-12 wks (**Figure 3A**) along with 4 other IHKA-injected animals (#5, 7, 8, 9) that were recorded at 10-12 wks only. A total of 473 seizures were analyzed and results are presented in **Figure 3B-H**. When all seizures during the two 2 wk-long periods (2-4 wks vs. 10-12 wks) were compared, 3 out of 7 animals (#1, 3, 12) that were recorded at both timepoints showed more seizures at 10-12 wks, an increase or progression (defined here as seizures that worsen with time), and the remaining 4 (#4, 6, 10, 11) showed a decrease (see **Figure 3B vs. Figure 2B**). We use the term progression conservatively as we cannot exclude the possibility that if a third timepoint was used the seizures that had increased from 2-4 to 10-12 wks might decrease eventually or vice-versa.

**Figure 3.**
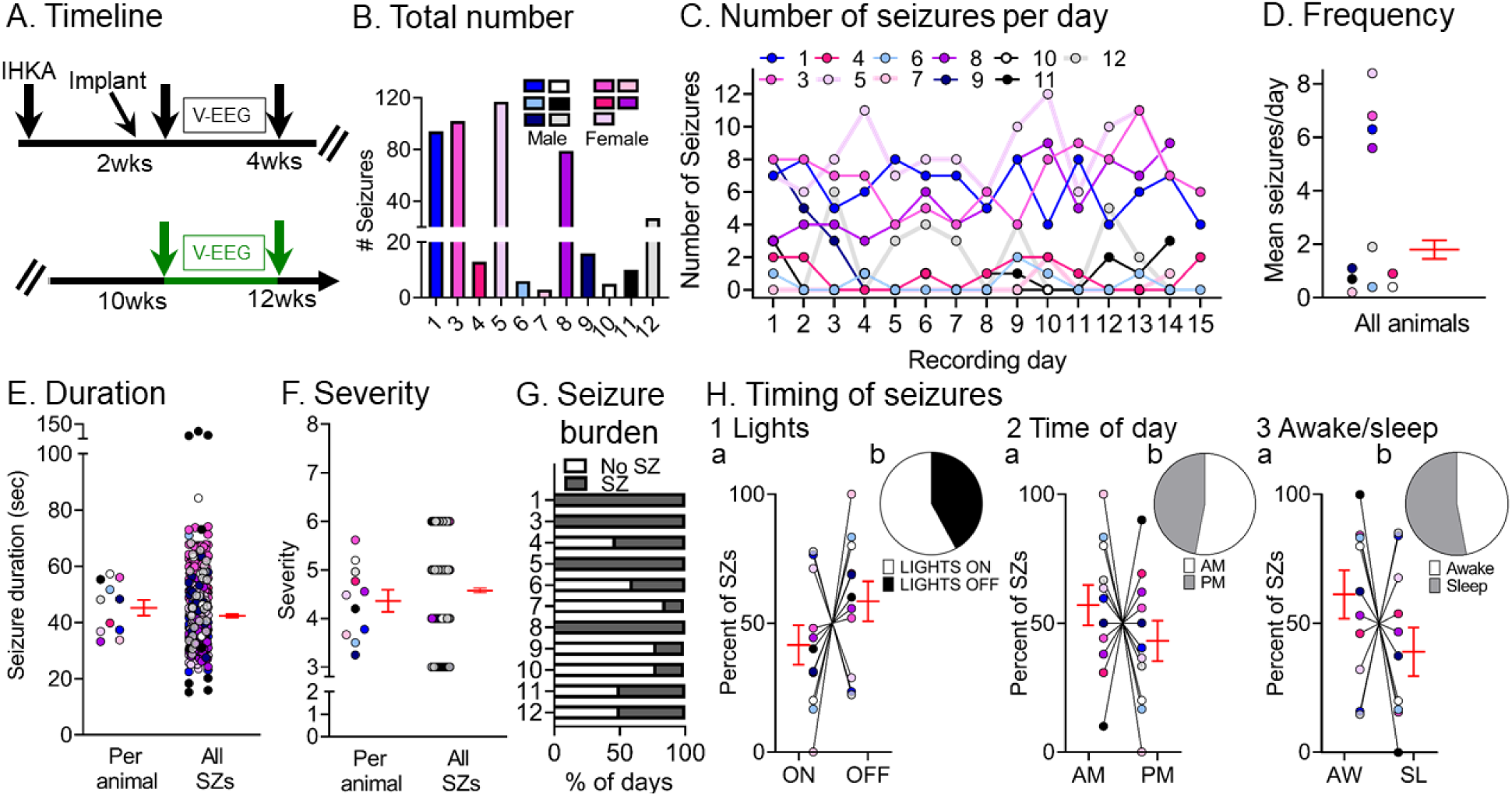
Quantification of chronic spontaneous convulsive seizures 10-12 wks post-IHKA. **(A)** Experimental timeline of the study. The 10-12 wk data were used for this figure (green arrows; total # seizures=473). Animals were recorded 2-4 wks post-IHKA and then again at 10-12 wks post-IHKA (5 male, 2 female). Additionally, 4 animals were recorded at 10-12 wks only (1 male, 3 female). **(B)** The total number of chronic convulsive seizures is shown per animal. Note that all animals showed convulsive seizures at 10 wks suggesting epilepsy persisted. **(C)** The number of convulsive seizures is plotted for each day of the recording period. As described in the legend of Fig. 2, each animal is a different line and color and the colors are consistent across figures. **(D)** Convulsive seizure frequency was calculated as the mean number of convulsive seizures per day for each animal. Data are presented in D-F and H as individual values and as mean ± SEM (red). **(E)** Convulsive seizure duration was calculated as the mean per animal (left) or the mean for all convulsive seizures (right; n=473 seizures). **(F)** Convulsive seizure severity was calculated as the mean per animal (left) or the mean of all convulsive seizures (right; n=473 seizures). **(G)** Convulsive seizure burden was defined as the percent of days spent with (SZ) or without (No SZ) convulsive seizures. **(H)** The percent of convulsive seizures is shown. either occurring during the light period or dark period of the light:dark cycle (Lights ON or OFF; H1), a.m. or p.m. (AM/PM; H2), and awake (AW) or sleep (SL) state (H3). The percentages were calculated as the mean per animal (H1a, H2a, H3a) or the mean of all seizures (H1b, H2b, H3b). There were no significant differences (paired t-tests; all p>0.05).

Similar to the 2-4 wk timepoint, convulsive seizures at the 10-12 wk timepoint were evident for most recording days (**Figure 3C**). There was a mean frequency of 2.9±0.9 (range 0.2-8.4) seizures per day per animal (**Figure 3D**). Convulsive seizures lasted for 45.2±2.8 sec (range 33.2-57.3) when the mean duration was calculated per animal and 42.4±0.6 sec (range 15.2-125.8) when all seizures were pooled (**Figure 3E**). These durations were significantly longer from those at 2-4 wks (**Supplemental Figure 2D**, paired t-test, *t*_crit_ = 3.14, p=0.02). Regarding severity, most seizures were severe because seizures scores were 4.3±0.2 (range 3-6) when the mean was calculated per animal and 4.6±0.04 (range 3-6) when all seizures were pooled (**Figure 3F**). These seizures scores were not significantly different from those at 2-4 wks (paired t-test, *t*_crit_ = 0.19, p=0.85). The mean percent of days with seizures (59.0±10.5%, range 14-100%) was not significantly different from the mean percent of days without seizures (40.7±10.4%, range 0-85%; Wilcoxon signed rank test, p=0.33), similar to the 2-4 wk timepoint.

For the animals with increased seizures at the late timepoint (#1, 3, 12), seizure burden was 83.3±16.7% and for the other mice (#6, 4, 10, 11) it was 41±7.2% (**Figure 3G**). When comparing the percent of seizures occurring during the time that lights were on vs. the time when lights were off, there were no significant differences (paired t-test, *t*_crit_ = 1.104, p=0.29; **Figure 3H1a, 3H1b**). There also were no significant differences between the percent of seizures during the a.m. vs. p.m. (paired t-test, *t*_crit_ = 0.886, p=0.39; **Figure 3H2a, 3H2b**). Finally, there was no difference in the percent of seizures occurring in awake vs. sleep states (paired t-test, *t*_crit_ = 1.184, p=0.26; **Figure 3H3a, 3H3b**).

A comparison of the data from 2-4 and 10-12 wks showed no significant differences in the total number (Wilcoxon signed rank test, p>0.99), frequency (Wilcoxon signed rank test, p>0.99), mean severity (paired t-test, *t*_crit_ =0.19, p=0.85) or days spent with seizures (unpaired t-test, *t*_crit_ = 0.14, p=0.88). Also, no differences were found in the percent of seizures occurring during lights on vs. lights off, a.m. vs. p.m. or awake vs. sleep between the 2 timepoints (**Supplemental Figure 3A**). The only difference was in seizure duration. Seizures at 2-4 wks were shorter than those recorded at 10-12 wks post-IHKA (paired t-test *t*_crit_ = 5.13, p=0.001). Therefore, taking seizure frequency into account, only 50% of mice increased seizure frequency between 2-4 and 10-12 wks, but if seizure duration is used as a measurement, the mice did exhibit worsening of their seizures between 2-4 and 10-12 wks post IHKA. The data are shown in more detail in Supplemental Figure 6A where seizures per day are plotted for consecutive days during the 2-4 wks post IHKA, and then 10-12 wks post IHKA. The data are also plotted for seizure duration (Supplemental Figure 6B).

### III. Chronic non-convulsive seizures occur after IHKA and are less frequent than convulsive seizures

Next, we analyzed non-convulsive seizures to determine whether they also are frequent post-IHKA. To that end, the same animals presented before in **Figures 1** and **2** were analyzed as shown in **Figure 4A**. An example of a non-convulsive seizure is shown in **Figure 4B1** and an expanded version of the same seizure is shown in **Figure 4B2**.

**Figure 4.**
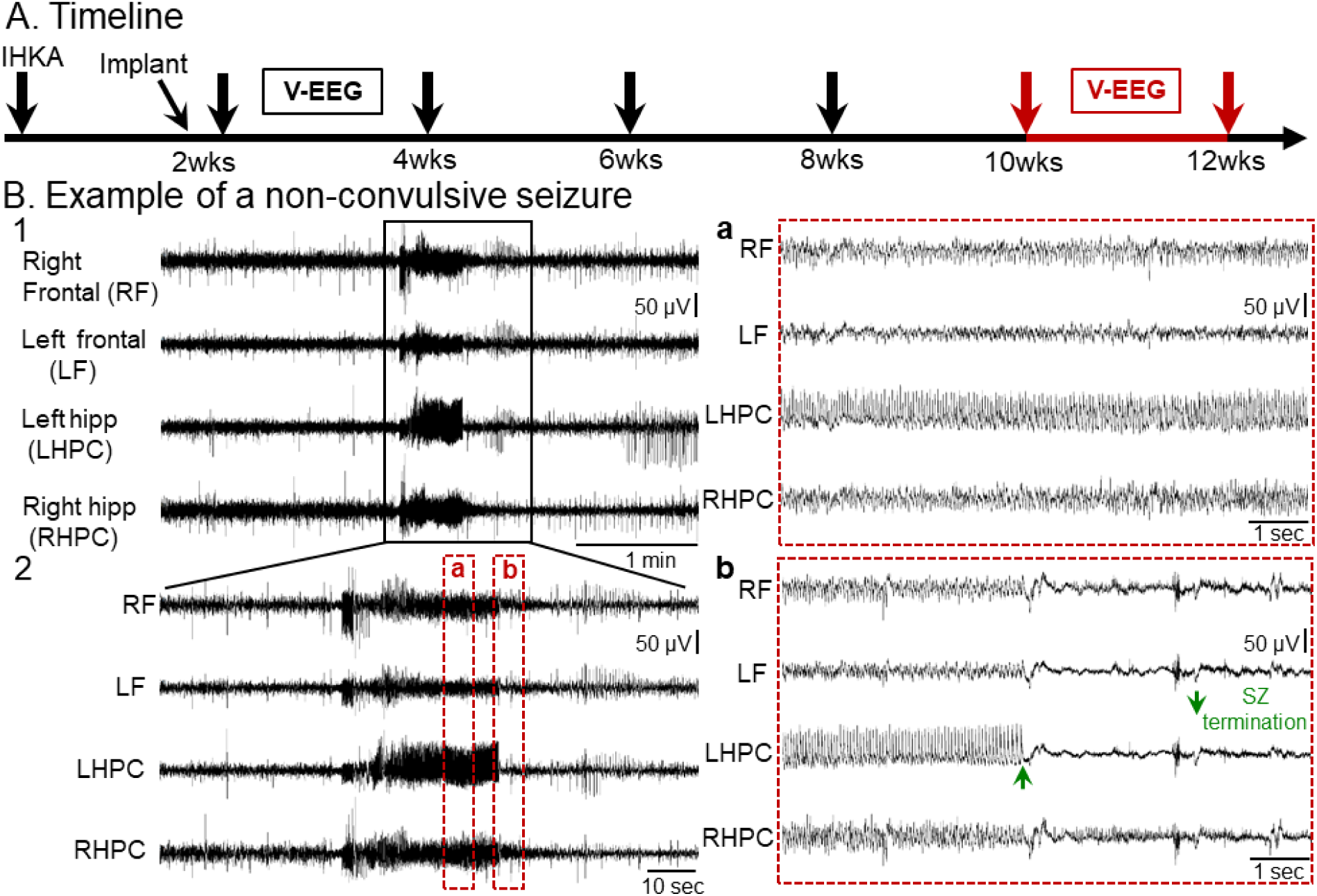
A spontaneous non-convulsive seizure recorded 10-12 wks post-IHKA. **(A)** Experimental timeline shows when the non-convulsive seizure in this figure was recorded, 10-12 wks post-IHKA (red arrows). For non-convulsive seizures, the mice were the same ones used to analyze convulsive seizures. **(B)** Representative example of a non-convulsive seizure recorded 10-12 wks post-IHKA. A 5 min-long EEG trace illustrating seizure activity in all 4 leads is shown in B1. The same seizure is depicted in a 2 min-long time window in B2 and further expanded in 10 sec-long epochs to better highlight electrographic activity over time. Inset a shows electrographic activity during the non-convulsive seizure. Inset b shows seizure termination. Note seizure termination was more distinct for some leads and the lead that was most distinct (green arrow in b, LHPC) varied from seizure to seizure.

The electrographic correlate of non-convulsive seizures consisted of a sudden increase in amplitude of rhythmic activity (>5 Hz) in all 4 leads. Insets in **Figure 4** show expanded EEG traces where the electrographic correlate of a non-convulsive seizure can be further appreciated. The non-convulsive seizure was characterized by trains of spikes of variable amplitude which appeared to occur synchronously in all 4 leads. Interestingly, post-ictal depression was not as pronounced as after a convulsive seizure (**see Figure 1B2b**).

We next quantified non-convulsive seizures at 2-4 wks (**Figure 5A**) and results are shown in **Figure 5B-H**. We found that both the total number and frequency of non-convulsive seizures were less frequent than convulsive seizures at 2-4 wks (paired t-test, *t*_crit_=4.16, p=0.03; Wilcoxon signed rank test, p=0.03, respectively) with no differences in mean duration (paired t-test, *t*_crit_=1.21, p=0.34); **Supplemental Figure 2D**). More detailed comparisons about lights on vs. off, a.m. vs. p.m. and awake vs. sleep are shown in **Supplemental Figure 3C** and **3D**.

**Figure 5.**
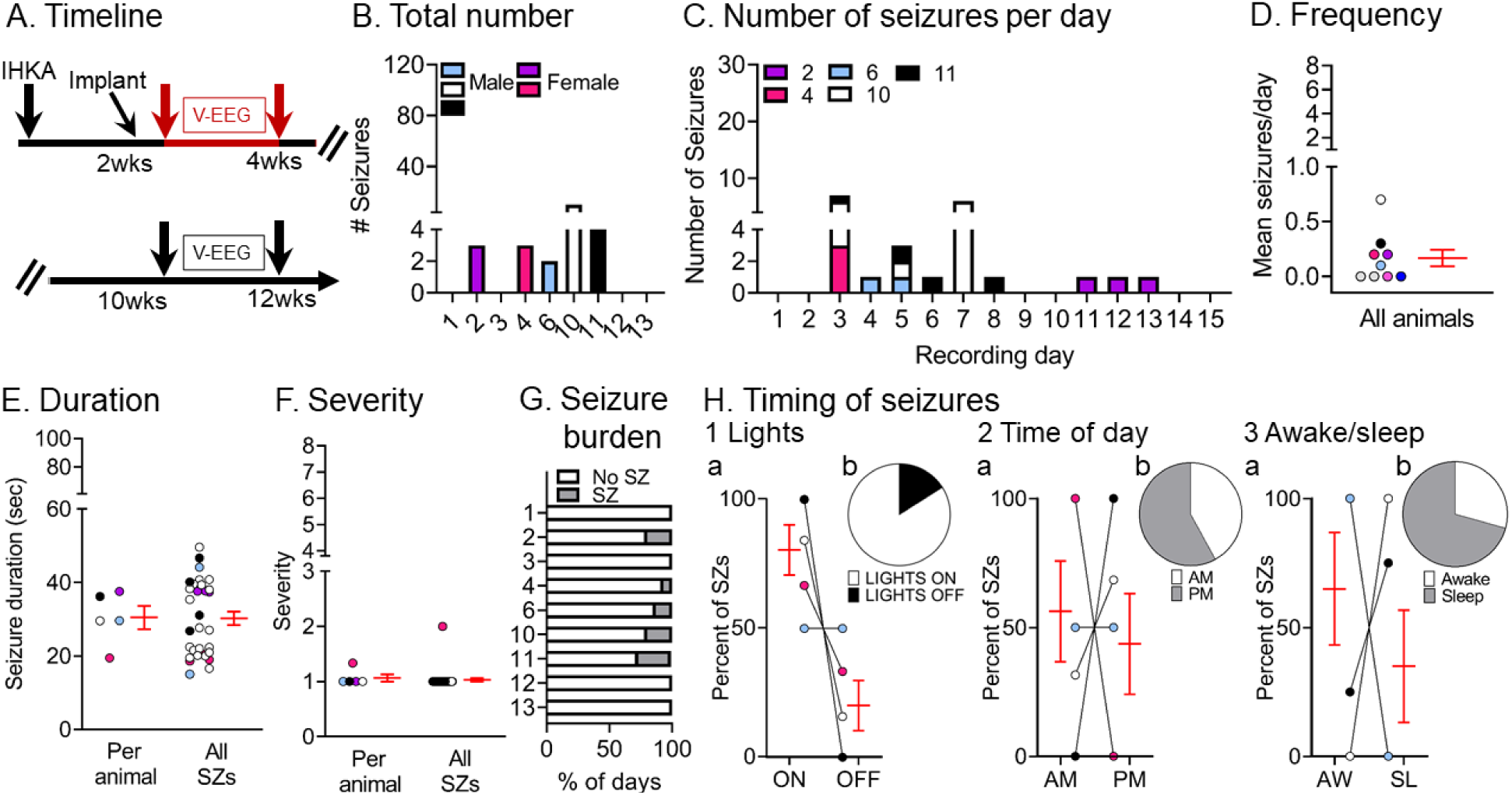
Quantification of chronic spontaneous non-convulsive seizures 2-4 wks post-IHKA. **(A)** Experimental timeline of the study. The seizures for this figure were recorded 2-4 wks after IHKA (red arrows; total # seizures=31). The mice were the same as those used for convulsive seizure measurements. **(B)** The total number of chronic non-convulsive seizures during the 2 wk-long recording period is shown per animal. Blue and pink shades represent males and females, respectively. **(C)** The number of non-convulsive seizures per day and per animal is shown for all recording days. **(D)** Non-convulsive seizure frequency was calculated as the mean number of non-convulsive seizures per day for each animal. In D-F and H, data are presented as individual values and as mean ± SEM (red). **(E)** Non-convulsive seizure duration was calculated as the mean per animal (left) or the mean of all non-convulsive seizures (right; n=31 seizures). **(F)** Non-convulsive seizure severity was calculated as the mean per animal (left) or the mean of all non-convulsive seizures (right; n=31 seizures). **(G)** Non-convulsive seizure burden was defined as the percent of days spent with (SZ) or without (No SZ) non-convulsive seizures in the 2-4 wk-long recording period. **(H)** The percent of non-convulsive seizures is shown, either occurring during the light period or dark period of the light:dark cycle (Lights ON or OFF; H1), a.m. or p.m. (AM/PM; H2), and awake (AW) or sleep (SL) state. Percentages are shown for the mean per animal (H1a, H2a, H3a) or the mean of all seizures (H1b, H2b, H3b). Note that in H2a there are 3 animals but the lines for 2 of the animals overlap since all of their seizures occurred during AM. In H3a, all seizures in all 3 animals occurred during the awake state (lines are overlapping). Statistical comparisons: H1a, not significant (Wilcoxon signed rank test; p>0.05); H2a, not significant (Wilcoxon signed rank test; p>0.05); H3a, not significant (Wilcoxon signed rank test; p>0.05).

**Figure 5C** shows the numbers of non-convulsive seizures for the different recording days, and it is evident that there could be few non-convulsive seizures. The mean frequency of non-convulsive seizures was 0.1±0.07 seizures per day (range 0-0.7; n=9 mice; **Figure 5D**) and the duration was 30.5±3.2 sec (range 19.4-37.6 sec) when the mean was calculated per animal and 30.2±1.8 sec (range 15.1-49.6 sec) when all seizures were pooled together (**Figure 5E**). The mean severity score was 1.1±0.06 (range 1-2) per animal and 1.0±0.03 (range 1-2) when all seizures were pooled (**Figure 5F**). The mean percent of days spent with vs. without seizures was 9.5±3.5% (range 0- 26%) and 90.3±3.5% (range 73-100%) respectively (**Figure 5G**) which was statistically significant (paired t-test, *t*_crit_=11.49, p<0.0001). No significant differences were found for lights on vs. off (Wilcoxon signed rank tests; all p>0.05; **Figure 5H1a, 5H1b**), a.m. vs. p.m. (Wilcoxon signed rank tests; all p>0.05; **Figure 5H2a, 5H2b**) or awake vs. sleep (Wilcoxon signed rank tests; all p>0.00; **Figure 5H3a, 5H3b**).

To determine whether non-convulsive seizures persisted with time, 4 out of 5 animals recorded at 2-4 wks were recorded again at 10-12 wks, as well as 3 more animals (78-3, 80-4, 81-1) that were recorded at 10-12 wks only (**Figure 6A**). Quantified non-convulsive seizures at 10-12 wks post-IHKA are shown in **Figure 6B-H**. Non-convulsive seizures at 2-4 wks and 10-12 wks were similar in number (Wilcoxon signed rank test, p=0.08; **Figure 6B, Supplemental Figure 2B**) and frequency (Wilcoxon signed rank test, p=0.11; **Figure 6C**, **Supplemental Figure 2C**). Seizure duration and severity are summarized in **Figures 6E** and **F** respectively and there was no significant difference from 2-4 wks (**Supplemental Figure 2D**). The percent of days spent with vs. without seizures did not change between timepoints (Wilcoxon signed rank test, p=0.25; **Figure 6G vs. Figure 5G**). There were also no significant differences in the proportion of seizures occurring during lights on vs. off (paired t-test, *t*_crit_=0.66, p=0.52), a.m. vs. p.m. (paired t-test, *t*_crit_=1.04, p=0.32) and awake vs. sleep states (paired t-test, *t*_crit_=0.26, p=0.79; **Figure 6H**), which did not differ with the 2-4 wk timepoint (**Supplemental Figure 3B**).

**Figure 6.**
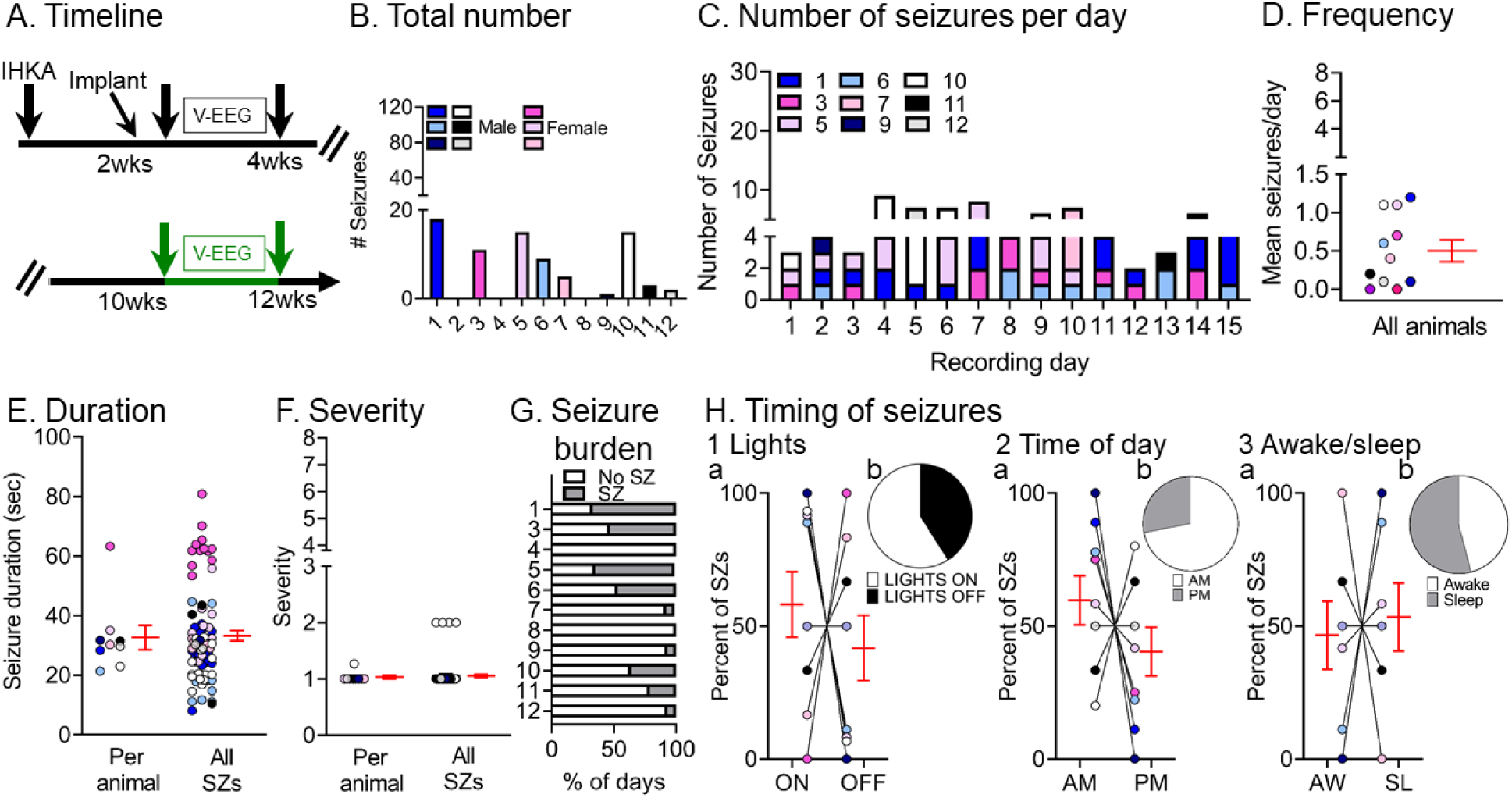
Quantification of chronic spontaneous non-convulsive seizures 10-12 wks post-IHKA. **(A)** Experimental timeline of the study. The seizures for this figure were recorded 10-12 wks after IHKA (green arrows; total # seizures=78). The mice were the same as those used for convulsive seizure measurements. **(B)** The total number of chronic non-convulsive seizures during the 2 wk-long recording period is shown per animal. Blue and pink shades represent males and females, respectively. **(C)** The number of non-convulsive seizures per day and per animal is shown for all recording days. **(D)** Non-convulsive seizure frequency was calculated as the mean number of non-convulsive seizures per day for each animal. In D-F and H, data are presented as individual values and as mean ± SEM (red). **(E)** Non-convulsive seizure duration was calculated as the mean per animal (left) or the mean of all non-convulsive seizures (right; n=78 seizures). **(F)** Non-convulsive seizure severity was calculated as the mean per animal (left) or the mean of all non-convulsive seizures (right; n=78 seizures). **(G)** Non-convulsive seizure burden was defined as the percent of days spent with (SZ) or without (No SZ) non-convulsive seizures in the 2 wks of continuous vEEG starting at 10 wks. **(H)** The percent of non-convulsive seizures is shown, either occurring during the light period or dark period of the light:dark cycle (Lights ON or OFF; H1), a.m. or p.m. (AM/PM; H2), and awake (AW) or sleep (SL) state (H3). The percentages were calculated as the mean per animal (H1a, H2a, H3a) or the mean of all seizures (H1b, H2b, H3b) respectively. They were no significant differences (paired t-tests; all p>0.05).

### IV. Different seizure onset patterns can be recorded and their prevalence changes with time

Recently several laboratories have suggested that seizures in rodent models of TLE can simulate human seizures (Velasco et al., 2000), and discussed several types of seizures based on their onset (Bragin et al., 1999b; Avoli et al., 2016). A common nomenclature refers to two types primarily: one type with low-voltage fast activity at the seizure onset (LVF seizure; (Bragin et al., 1999b) and another type with a hypersynchronous onset (HYP; (Bragin et al., 1999b).

The seizures we recorded showed onset patterns that remarkably, almost always fit the description of LVF and HYP seizures. The LVF pattern was characterized by a “sentinel” spike and by brief suppression of background EEG followed by rhythmic activity (**Figure 7A, Supplemental Figure 7**). The HYP pattern was characterized by a progressive increase in spike frequency (2-5 Hz) that eventually escalated to the point where a seizure was clear (**Figure 7B, Supplemental Figure 8**). However, there were some seizures that could not fit into the two categories, which we classified as “unclear” onset.

**Figure 7.**
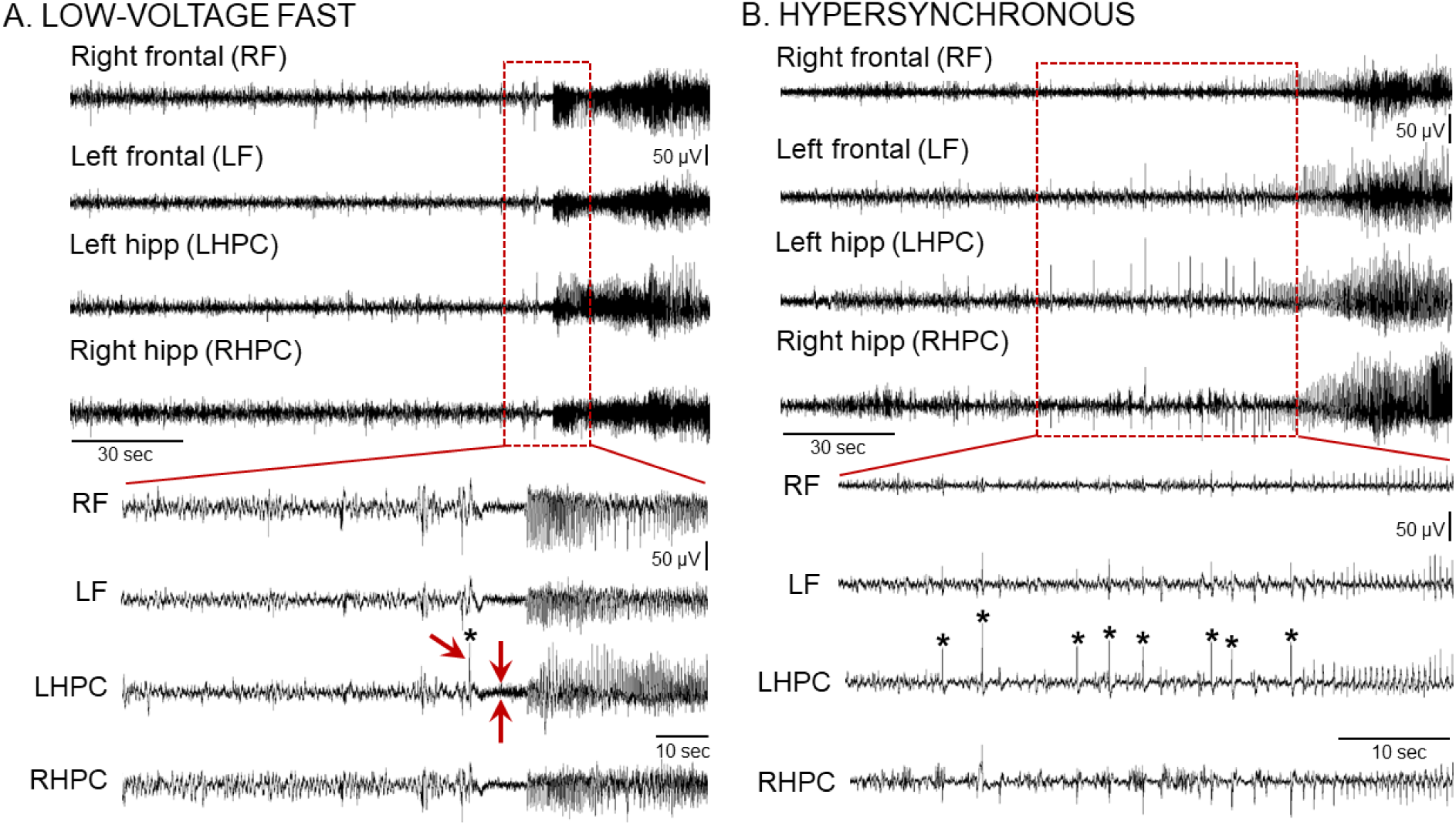
Chronic IHKA seizures show different seizure onset patterns. **(A)** Representative example of a low-voltage fast (LVF) onset seizure recorded during the 10-12 wk recording session post-IHKA. A 30 sec-long EEG trace (Top) and an expanded 10 sec-long epoch (Bottom) taken from the seizure onset are shown. Note that the LVF pattern starts with a sentinel spike (asterisk and single red arrow in the expanded trace at the Bottom) followed by brief suppression of the background EEG (2 red arrows pointing at each side of the EEG suppression) and subsequent series of spikes. Additional examples of LVF seizures are shown in **Supplemental Figure 7**. **(B)** Representative example of a hypersynchronous (HYP) onset seizure recorded during the 2-4 wk recording period post-IHKA. A 2 min-long EEG trace (Top) and an expanded 30 sec-long epoch (Bottom) taken from the seizure onset are shown. Note that the HYP patterns start with a series of spikes (asterisks) followed by spikes at increased frequency. Additional examples of HYP seizures are shown in **Supplemental Figure 8**.

Next, we quantified the different seizure onset patterns to understand if one of the two patterns was more prominent (**Figure 8**). For this analysis, we analyzed a total of 800 seizures (all data from each timepoint were pooled). Remarkably, almost all seizures either fit the LVF or HYP pattern (**Figure 8B**). There were no significant differences between LVF and HYP seizures in the total number (Wilcoxon sighed rank test, p=0.072; **Figure 8B**) or frequency (Wilcoxon signed rank test, p=0.77; **Figure 8C**). When seizure durations were measured for each animal, the means were not different (paired t-test, p>0.05; **Figure 8D1**). However, when all seizure durations were pooled, LVF seizures lasted significantly longer than HYP seizures (Wilcoxon signed rank test, p<0.0001; **Figure 8D2**). The severity was not different when it was measured for each animal (paired t-test, p>0.05; **Figure 8E1**), but LVF seizures were significantly more severe when all seizures were pooled (Wilcoxon signed rank test, p=0.02; **Figure 8E2**), probably because the sample size for severity was animals and the sample size for all seizures was much larger.

**Figure 8.**
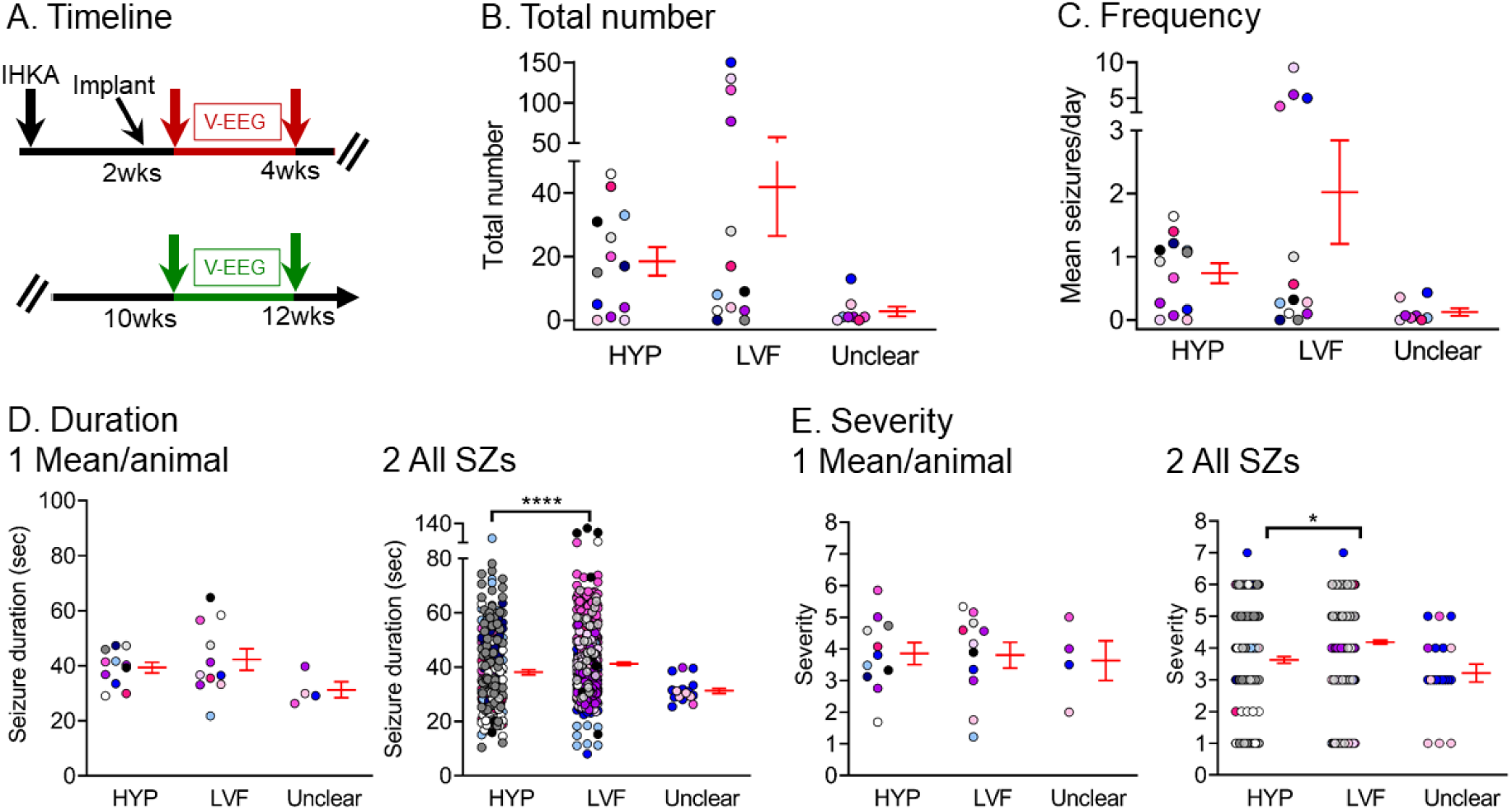
Quantification of low-voltage fast and hypersynchronous onset seizures. **(A)** Experimental timeline of the study. The data used for this figure were pooled from the 2-4 wk (red arrows) and 10-12 wk timepoints (green arrows) and a total of 800 seizures were included in the analyses. Seizures were distinguished as HYP, LVF, or seizure onset type was unclear. There were 8 mice (2 males, blue, recorded at both 2-4 and 10-12 wks; 6 females, pink, where 2 were recorded at both times, and 4 were recorded only at one of the times). **(B)** The total number of seizures is shown. There was no significant difference between HYP and LVF seizures (Wilcoxon signed rank test, p=0.72). In B-E, data are presented as individual values and as mean ± SEM (red). **(C)** Same as B but the frequency of seizures is plotted. There was no significant difference between HYP and LVF seizures (Wilcoxon signed rank test, p=0.77). **(D)** Same as B but seizure duration is shown. Seizure duration was calculated as an average per animal (D1), or all seizures were pooled (D2). LVF seizures lasted significantly longer than HYP seizures when all seizures were pooled (D2; Wilcoxon signed rank test, p<0.0001) and not when duration was calculated as an average per animal (D1; paired t-test, p>0.05). **(E)** Same as B but seizure severity is shown. Seizure severity was calculated as an average per animal (E1) and for all seizures (E2). LVF seizures were significantly more severe than HYP seizures when all seizures were pooled (E2; Wilcoxon signed rank test, p=0.02). In other words, E1 was not significant but E2 was, and that result is likely to be due to the large number of seizures in E2 compared to animals in E1.

We next evaluated if there was a preference towards a particular seizure onset pattern between the 2 different timepoints, 2-4 or 10-12 wks. Results are presented in **Figure 9**. HYP seizures were more frequent early whereas LVF seizures were more frequent late (HYP: paired t-test, *t*_crit_=5.63, p=0.0013; LVF: Wilcoxon signed rank test, p=0.04; **Figure 9B**). Interestingly, when the percent of HYP and LVF seizures were analyzed instead of the total number, HYP and LVF seizures were significantly different between timepoints with more HYP seizures dominating the earlier timepoint and more LVF seizures dominating the later timepoint (Fisher’s exact test, p<0.0001; **Figure 9C**).

**Figure 9.**
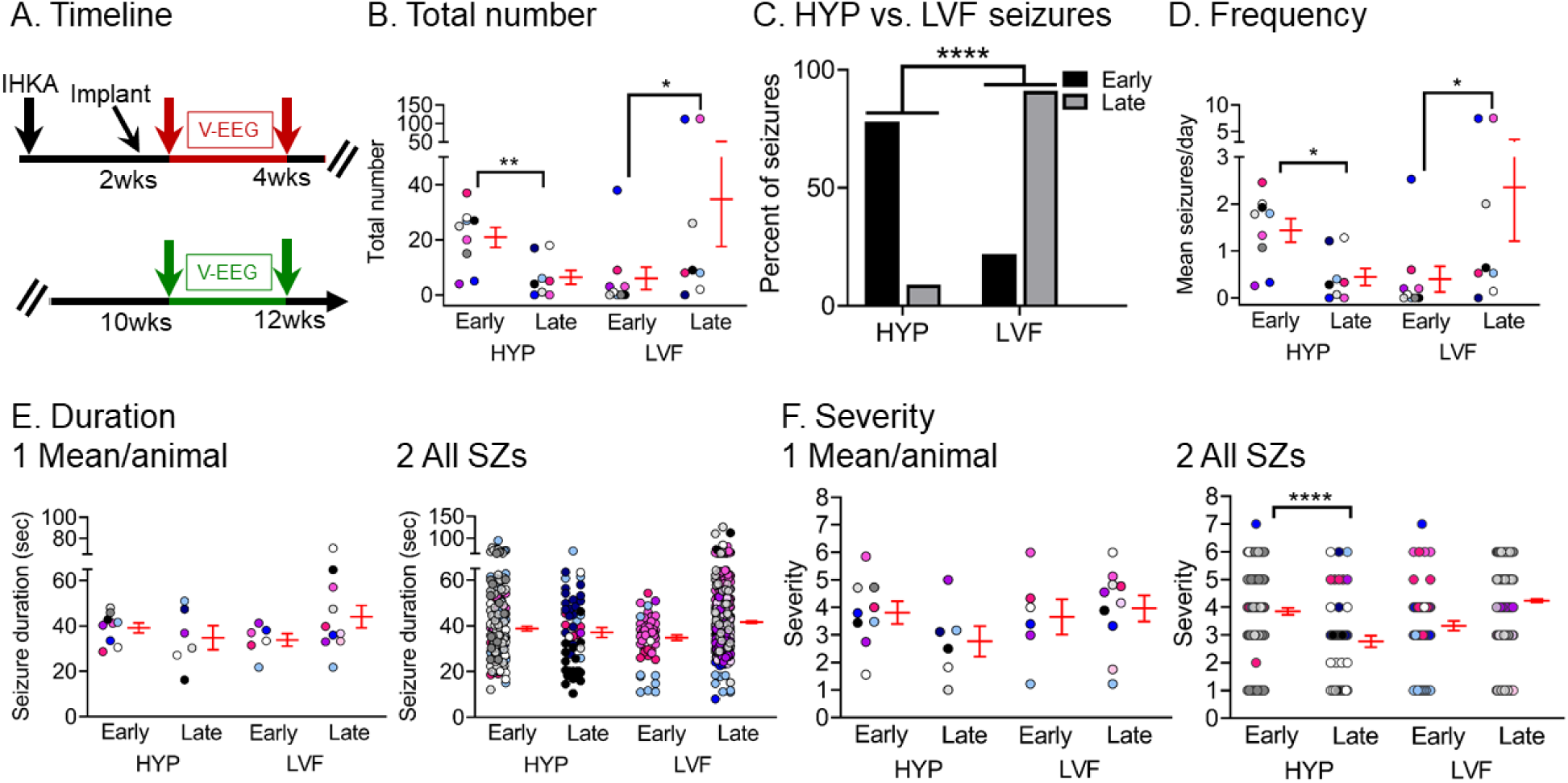
Quantification of different seizure onset patterns between timepoints. **(A)** Experimental timeline of the study. The data used for this figure were both from the 2-4 wk (red arrows) and 10-12 wk timepoint (green arrows). Seizures were distinguished as HYP or LVF. Seizures that were unclear in onset are not included in this figure because they were relatively rare (see Figure 8). **(B)** The total number of chronic HYP and LVF seizures recorded at the early (2-4 wks) and late (10-12 wks) timepoints are shown. HYP seizures were more frequent early whereas LVF seizures were more frequent late (HYP: paired t-test, *t*_crit_=5.63, p=0.0013; LVF: Wilcoxon signed rank test, p=0.04; Figure 9B). In B and D-F, data are presented as individual values and as mean ± SEM (red). **(C)** The percent of HYP vs. LVF seizures are shown for early and late timepoints. The percentage for each seizure category is shown for each timepoint. HYP seizures dominated early timepoints whereas LVF seizures dominated late timepoints (Fisher’s exact test, p<0.0001). **(D)** HYP and LVF seizure frequency was calculated as the mean number of seizures per day for each timepoint (early, late). HYP seizures were more frequent early vs. late (Wilcoxon signed rank test, p=0.01), whereas LVF seizures were more frequent late vs. early (Wilcoxon signed rank test, p=0.01). **(E)** HYP and LVF seizure duration was calculated as the mean number of seizures per day for each animal at each timepoint (early, late; E1). No significant differences were found (Wilcoxon signed rank test, p>0.05). (E2) Same as in E1 but all HYP and LVF seizures were pooled and are presented according to timepoint and type (HYP, LVF). No significant differences were found (Wilcoxon signed rank test, p>0.05). **(F)** HYP and LVF seizure severity was calculated as the mean per animal at each timepoint (early, late; F1). (F2) Same as in F1 but all seizures were pooled and are presented according to timepoint and type (HYP, LVF). HYP early seizures were less severe than HYP late seizures (Wilcoxon signed rank test, p<0.0001).

The same result was found by analysis of the frequency of LVF or HYP seizures (**Figure 9D**), HYP seizures were more frequent at 2-4 wks and LVF seizures were more frequent at 10-12 wks (HYP: Wilcoxon signed rank test, p=0.01, LVF: Wilcoxon signed rank test, p=0.01; **Figure 9B**). No significant differences were found in durations (Wilcoxon signed rank tests, p>0.05; **Figure 9E1, 9E2**).

However, HYP early seizures were less severe than HYP late seizures when all seizures were pooled (Wilcoxon signed rank test, p<0.0001; **Figure 9F2**).

Similar results were obtained when convulsive seizures were analyzed only (**Supplemental Figure 4**). A more detailed comparison of convulsive seizure onset patterns among animals that showed an increase (‘progressed’) vs. a decrease in seizures (‘non-progressed’) between timepoints is shown in **Supplemental Figure 5**.

### V. HFOs (>250 Hz) after IHKA are frequent at the site of IHKA injection, during seizures, and also occur interictally

Additional analysis of the EEG traces revealed the presence of HFOs in the frequency range of 250-500 Hz. HFOs were frequent during slow wave sleep, in line with previously published observations (Staba et al., 2004). Examples of interictal HFOs are shown in **Figure 10B** and their spectral properties are shown in **Figure 10D1, D2**.

**Figure 10.**
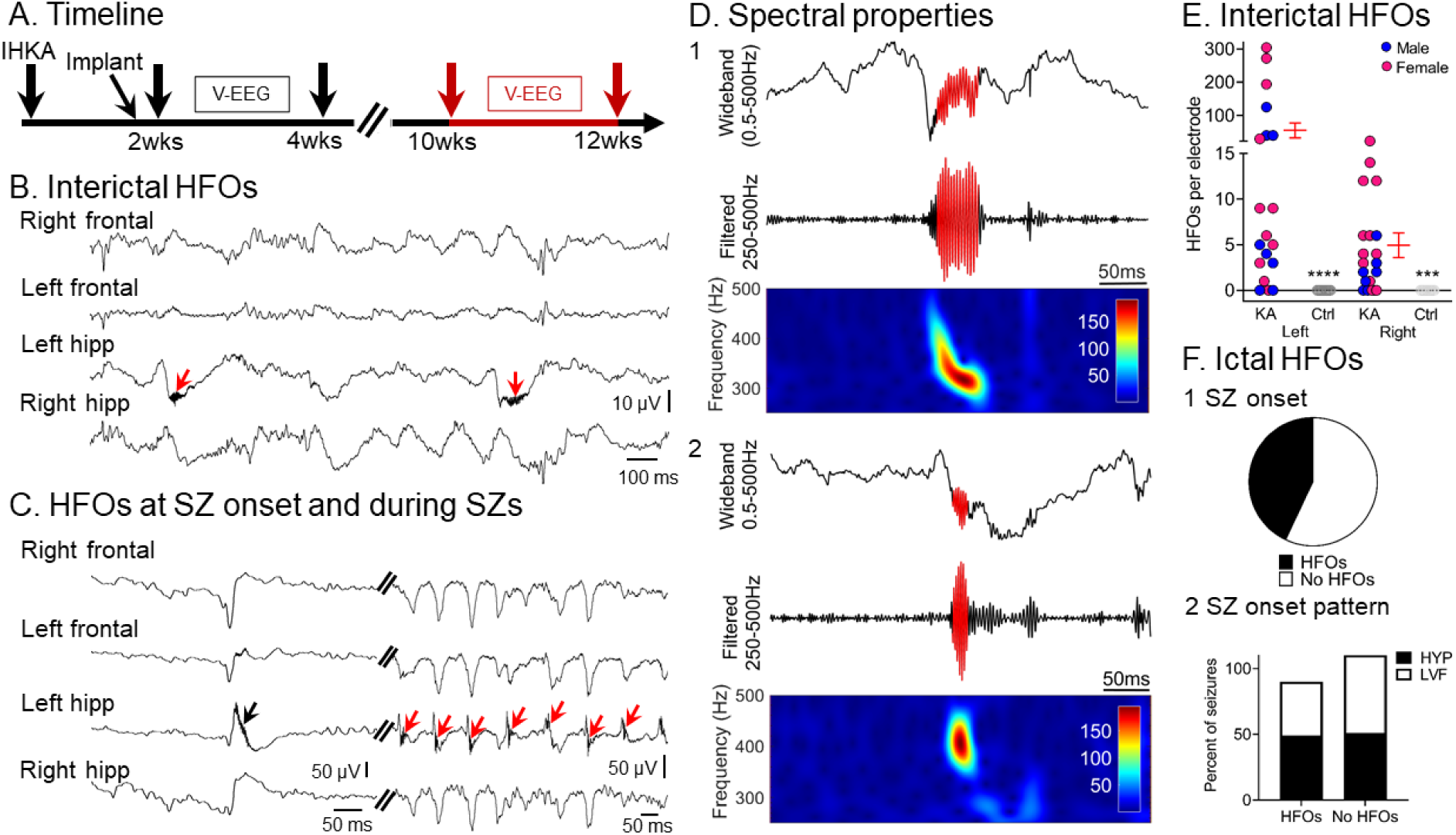
HFOs (>250 Hz) are frequent interictally, before and during seizures, and are recorded primarily from the IHKA injection site. **(A)** Experimental timeline of the study. The data used for this figure come from the 10-12 wk timepoint (red arrows). **(B)** Representative example of interictal HFOs recorded from the left hemisphere. Note the presence of HFOs (red arrows) in the left hippocampal lead where they were in the trough of slow waves. Note that interictal HFOs were not always in the trough of slow waves, however. **(C)** An example of an HFO recorded at the sentinel spike of an LVF seizure (black arrow) as well as during the seizure (red arrows). In this instance, HFOs were primarily recorded from the left hippocampal lead but this was not always the case. **(D)** Two examples of HFOs (D1, D2). For both D1 and D2, the top is wideband recording (0.5-500 Hz) showing coupling of HFOs to the trough of slow waves. The centers are the filtered traces (250-500 Hz) and the bottom shows spectral properties of HFOs with frequencies >250 Hz in the 250-500 Hz time-frequency domain. **(E)** Quantification of HFOs (events/min) based on either the left (hippocampus or cortex) or right (hippocampus or cortex) electrodes for IHKA and Saline-injected animals. HFOs were more frequent in the left hemisphere compared to the right (Wilcoxon signed rank test, p=0.01). HFOs were not detected in controls. Comparisons of IHKA- and Saline-injected mice showed increased HFOs in IHKA-injected mice for both left and right electrodes (Left: Mann-Whitney U-test, U = 18, p<0.0001, Right: Mann-Whitney U-test, U = 30, p<0.0001). Blue and pink represent males and females, respectively. Data are presented as individual values and as mean ± SEM (red). **(F)** The percent of seizures that showed HFOs at their onset (F1). The percentage of HYP vs. LVF seizures that showed HFOs at seizure onset was not statistically different (Fisher’s exact test, p>0.05; F2). The percentage of HYP and LVF seizures for each category (HFOs, no HFOs) was calculated for the total number of seizures for each seizure type (HYP or LVF).

HFOs also appeared frequently at seizure onset and during seizures. An example of HFOs recorded at seizure onset and during an LVF seizure is shown in **Figure 10C**. Forty-three percent of all recorded seizures showed HFOs at seizure onset (**Figure 10F1**). When we looked at the percentage of HYP vs. LVF seizures that showed and did not show HFOs at their onset we did not find a significant difference (Fisher’s exact test, p>0.05; **Figure 10F2**). Thus, a similar percent of HYP vs. LVF seizures seemed to be associated with HFOs at seizure onset (Fisher’s exact test, p>0.05; **Figure 10F2**). The percentage of HYP and LVF seizures for each category (HFOs, no HFOs) was calculated for the total number of seizures for each seizure type (HYP or LVF).

HFOs were more frequent in the left compared to the right hemisphere (Wilcoxon signed rank test, p=0.01; Figure 10E). Importantly, we found that interictal HFOs occurred in both sexes. The mean number of HFOs recorded from the left hemisphere (hippocampus or cortex) in the males was 27.1±15.2 and in the females was 75.7±35.9. However, sex differences were not significant (Mann-Whitney U-test, U = 34, p=0.42).

None of the Saline-injected controls (n=6) showed HFOs (Supplemental Table 1).

### VI. Hippocampal damage to the ipsilateral hippocampus is consistent with MTS and correlates with SE severity but not chronic seizure burden

To determine post-IHKA neuropathological outcomes, animals were sacrificed at 12 wks post-IHKA (**Figure 11A**) and coronal brain sections located adjacent and more posterior to the IHKA injection site were stained with cresyl violet (**Figure 11**). No damage was detected in Saline-injected controls (**Figure 11B**) Neuronal loss was primarily in the ipsilateral hippocampus (**Figure 11C**) and included loss in the hilus and pyramidal cell layers (PCL) of CA3 and CA1. Damage to the hippocampal pyramidal cells near the IHKA injection site, and posterior to it, was estimated by measuring the length of the PCL where neurons survived. For this purpose, measurements were made from, the border of CA1 with the subiculum through the entirety of CA1, CA3, and CA3, ending at the border of CA3 with the hilus (**Supplemental Figure 1**). There was a significant reduction in PCL length in the dorsal areas near the KA injection site compared to controls (Mann-Whitney U-test, U = 0, p=0.0008, n=17; **Figure 11D1**). There also was significant damage in the PCL posterior to the KA injection site (**Figure 11E1**).

**Figure 11.**
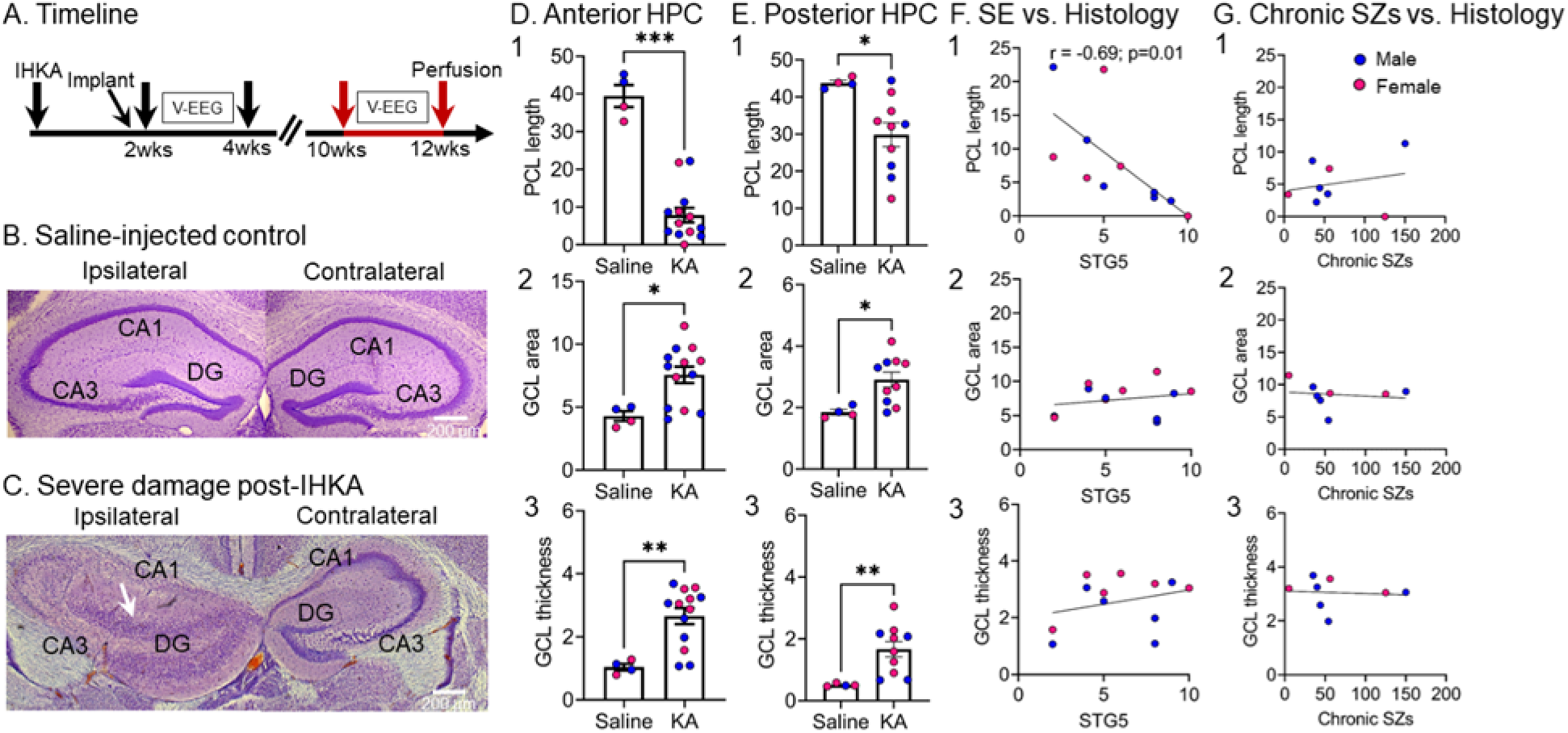
Quantification of hippocampal damage at 12 wks post-IHKA. **(A)** Experimental timeline of the study indicating the time animals were perfused at 12 wks post-IHKA (red arrows). Neuronal damage was quantified based on cresyl violet-stained coronal sections that were adjacent (-2 mm A-P, ± 1.25 mm M-L, 1.6 mm D-V) and more posterior (-2.5 mm A-P, ± 1.25 mm M-L, 1.6 mm D-V) to the IHKA injection site corresponding approximately to plate 47 and 51 of a standard mouse stereotaxic atlas (Franklin and Paxinos, 1997). **(B)** An example of a Saline-injected control showing that there is no obvious hippocampal damage. Calibration bar: 200 µm. **(C)** An example of severe hippocampal damage adjacent to the IHKA injection site. Note extensive neuronal loss in all pyramidal layers of the left hippocampus. Also, note extensive granule cell dispersion (white arrow). Calibration bar: 200 µm. **(D)** Measurements of the IHKA-injected hippocampus adjacent to the IHKA injection site are shown. Neuronal damage in the pyramidal cell layers was quantified by the length of the pyramidal cell layer (PCL) which did not exhibit neuronal loss. Note the reduced PCL length in IHKA vs. Saline-injected animals (Mann-Whitney U-test, U = 0, p=0.0008; D1). Granule cell layer (GCL) area was used as a reflection of granule cell dispersion with larger GCL area in IHKA vs. Saline-injected animals (Mann-Whitney U-test, U = 7, p=0.031; D2) The width of the GCL was used as another measure of granule cell dispersion, confirming that the GCL is wider in IHKA vs. Saline-injected animals (Mann-Whitney U-test, U = 4, p=0.01; D3). **(E)** Same as in D, but measurements were taken for sections that were more posterior to the IHKA injection site. Note significant reduction in PCL length (Mann-Whitney U-test, U = 3, p=0.01; E1), increased GCL area (Mann-Whitney U-test, U = 3, p=0.01; E2) and thickness (Mann-Whitney U-test, U = 0, p=0.002; E3) compared to Saline-injected controls. **(F)** There was a significant correlation between the number of stage 5 seizures during IHKA-induced SE and PCL length (r=-0.69, p=0.01; F1), but not GCL area (r=0.22, p=0.48; F2) and not GCL thickness (r=0.29, p=0.35; F3). All IHKA-injected animals were included in the analyses. **(G)** There was no significant correlation between the total number of chronic seizures (at both timepoints; 2-4 and 10-12 wks) and PCL length (r=0.23, p=0.57; G1), GCL area (r=-0.15, p=0.71; G2) or GCL thickness (r=-0.08, p=0.84; G3) at 12 wks post-IHKA. For the total number of chronic seizures, seizures were pooled for the two timepoints and only those animals with recordings at both timepoints are included.

To quantify granule cell dispersion, the granule cell layer (GCL) area was measured, as well as the thickness (**Supplemental Figure 1**). There was a greater GCL area in IHKA-treated animals compared to Saline-treated controls (Mann-Whitney U-test, U = 7, p=0.031; **Figure 11D2**), suggesting granule cell dispersion after IHKA. In addition, there was increased GCL thickness in IHKA-treated animals vs. Saline-injected controls (Mann-Whitney U-test, U = 4, p=0.01; **Figure 11D3**). Same quantifications at a more posterior to IHKA brain level confirmed significant PCL reduction (Mann-Whitney U-test, U = 3, p=0.01; **Figure 11E1**), increased GCL area (Mann-Whitney U-test, U = 3, p=0.01; **Figure 11E2**) and thickness (Mann-Whitney U-test, U = 0, p=0.002; **Figure 11E3**) compared to controls suggesting hippocampal damage extended posterior to the IHKA injection site.

To address the relationship between hippocampal damage and seizures during SE, PCL length, GCL area, and GCL thickness were plotted relative to the number of stage 5 seizures during SE (**Figure 11E, F**). PCL length was significantly correlated with the number of stage 5 seizures, such that PCL length was shorter when mice had more stage 5 seizures (r=-0.69, p=0.01; **Figure 11E1**), indicating the more severe seizures during SE were associated with more neuronal loss in the PCL. The idea that more neuronal damage occurs when SE is more severe is consistent with prior studies of SE (Dingledine et al., 2014). GCL area was not correlated significantly with the number of stage 5 seizures during SE (r=0.22, p=0.48; **Figure 11E2**) and GCL thickness was not either (r=0.29, p=0.35; **Figure 11E3).** These data are consistent with the idea that GCL dispersion may not be a result of the severity of SE but additional factors (Haas et al., 2002; Kobow et al., 2009).

To address the relationship between hippocampal damage and the number of chronic seizures, the PCL length, GCL area, or GCL thickness were plotted relative to the total number of chronic seizures at both timepoints (2-4 and 10-12 wks) post-IHKA (**Figure 11F**). The correlations were not significant (PCL length: r=0.23, p=0.57, **Figure 11F1;** GCL area: r=-0.15, p=0.71, **Figure 11F2**; GCL thickness: r=-0.08, p=0.84, **Figure 11F3)**. The reasons that PCL length did not correlate with chronic seizure burden is consistent with the idea that PCL neuronal loss is primarily due to SE-associated excitotoxicity (Dingledine et al., 2014). However, one would expect more damage to lead to more chronic seizures. The reason that estimates of GCL dispersion were not correlated with chronic seizure burden may be due to the idea that it arises during epileptogenesis but not as a result of chronic seizures (Haas et al., 2002; Kobow et al., 2009).

## IV. DISCUSSION

### A. Summary of main findings

The results suggest that our use of IHKA in mice results in robust spontaneous convulsive seizures. The seizures are robust because of their severity and duration. Also, animals had frequent convulsive seizures that continued for over 2 wks. In addition, these seizures were robust because the electrographic correlates showed high amplitude, long-lasting rhythmic activity in each of our 4 leads. These seizures ended with postictal suppression of the EEG, which is another characteristic that supports the description of the seizures as robust. These seizures were evident 2-4 wks after IHKA and at 10-12 wks, but some animals showed an increase in frequency and others showed a decrease, suggesting variability but robust epilepsy, nevertheless. Notably, seizure duration consistently increased with time even if frequency did not, suggesting progressively worse epilepsy depending on the type of measurement of seizures. Non-convulsive seizures were also evident but were less frequent than convulsive seizures. Interestingly, seizures did not appear to be focal, meaning they were not confined to one electrode (such as the site where IHKA was infused).

Convulsive seizures showed seizure onset patterns other investigators have found in their rat models of epilepsy (Bragin et al., 1999b; Bragin et al., 2005; Lévesque et al., 2012), as well as humans with TLE (Velasco et al., 2000; Perucca et al., 2014; Gnatkovsky et al., 2019; Saggio et al., 2020). HFOs were frequent not only at the site of IHKA injection but the ipsilateral cortical site, suggesting a wider epileptogenic network beyond the IHKA injection site. Although HFOs were observed in the contralateral hemisphere, they were relatively rare. Nevertheless, diverse sites of HFOs suggest caution before assuming they are the site of the seizure focus, consistent with a previous study showing HFOs outside the IHKA focus in mice (Sheybani et al., 2018). However, the rapid generalization of the seizures we recorded did not permit us to ascertain where the focus was, and there could have been more than one focus.

Histopathological findings were consistent with prior reports about the IHKA model (Bouilleret et al., 1999), including extensive hippocampal neuronal loss in the area of IHKA injection, like MTS. Although other types of quantification are common, such as cell counts, our conclusion that there was extensive pyramidal cell loss would have been clear with almost any measurement, in our view. For this reason, and to avoid the uncertainty of cell counts in the packed cell layers, we did not count cells. We mention in the Results that we consider the measurement of PCL damage an estimate only. Besides neuronal loss, granule cell dispersion was also evident in our animals, as reported before (Bouilleret et al., 1999). Far less disruption of the contralateral hippocampus is also consistent with prior reports (Bouilleret et al., 1999).

### B. IHKA produces robust chronic epilepsy

Several electrophysiologic elements of chronic convulsive seizures reported here are distinct from the very frequent short-lasting epileptiform abnormalities reported before (Kim et al., 2018; Sandau et al., 2019; Lai et al., 2020). One element is seizure duration. We provide evidence of prolonged convulsive seizures similar to seizure durations reported in human TLE (Balish et al., 1991) and data from an Epilepsy Monitoring Unit (Jenssen et al., 2006). The longer-lasting seizures contrast with the 3-7 sec epileptiform abnormalities reported by some past studies of IHKA in mice (Kim et al., 2018; Sandau et al., 2019; Lai et al., 2020). In addition, our results differ from what appears to be continuous spiking at the site of IHKA injection indicated by some investigators (Sandau et al., 2019).

Another aspect of the seizures we recorded was the presence of complex rhythmic activity during a seizure. Thus, seizures we recorded were more complex electrographically than some trains of spikes shown by others (Kim et al., 2018; Sandau et al., 2019; Lai et al., 2020). For instance, our seizures showed fast and slow spikes as well as repetitive spikes and waves at many different frequencies. These complex EEG patterns were found for both convulsive and non-convulsive seizures. Finally, the convulsive seizures we recorded were typically followed by prolonged post-ictal suppression and this was not clear in studies by others (Kim et al., 2018; Sandau et al., 2019; Lai et al., 2020).

Another aspect of the EEG that we found was a characteristic of seizures (both convulsive and non-convulsive) was that they were recorded at 4 sites, both hippocampi and two cortical locations. In the past, many studies of the IHKA model in mice did not use as many recording sites so less information was available (Riban et al., 2002; Zeidler et al., 2018). On the basis of their recording sites and the lack of convulsive behavior, conclusions were sometimes made that the model showed focal seizures primarily (Riban et al., 2002). An important point about our recordings is that we found synchronized activity even from the start of the seizure. This was especially clear for LVF seizures because at the onset there is a sentinel spike.

Regarding the site of the seizure focus, based on the findings from sentinel spikes, one might argue that seizure onset is where the sentinel spike is largest, the hippocampus. This interpretation seems logical since it was the site of IHKA injection. However, the subdural screws for cortical recordings had less resolution than depth electrodes in the hippocampus. In addition, it has been suggested that the origin of LVF seizures is extrahippocampal (Velasco et al., 2000).

Chronic convulsive seizures were detected both at 2-4 and 10-12 wks, another argument that chronic epilepsy was robust because convulsive seizures continued to occur. The fact that seizures persisted is important to be noted as there is little quantified longitudinal data of chronic seizure outcomes beyond 2 months post-IHKA in mice (Henshall, 2017). However,1 or 2 motor seizures a wk and seizures at 8 months were mentioned in the text of the study of Bouilleret et al. (Bouilleret et al., 1999).

We found that at 10-12 wks after IHKA, 50% of animals increased their seizure frequency and the rest decreased their seizure frequency compared to the 2-4 wk timepoint. None of the animals showed an absence of seizures at 10-12 wks. The variability in seizure frequency is important because it is consistent with human TLE. Thus, seizure diaries of patients with TLE suggest that some individuals may experience an increase in seizures with time whereas others report a decrease (Bauer and Burr, 2001). However, it is important to also note that patients were taking ASDs and our animals were not. Also, seizure diaries are not as accurate as continuous vEEG. The variable seizure frequencies we found have important implications for preclinical drug testing, because testing is usually for only one time period, and that lasts only 2 wks. Our data suggest that a 2 wk-long testing period may not predict seizure frequency throughout the lifespan. Testing at different timepoints during chronic epilepsy has been stressed by human studies already (Zimmermann and Trinka, 2020; Thomson et al., 2021).

One study of IHKA is important to note because the investigators did study progression of the epilepsy and no clear progression was found (Welzel et al., 2019). Those data are consistent with our findings in that we found progression only in half of the animals.

### C. Chronic IHKA seizures show different seizure onset patterns

The original studies of IHKA in mice have suggested no changes in seizure patterns with time (Bouilleret et al., 1999). However, a limitation to assess this might be related to the fact that convulsive seizures were rare, or discontinuous vEEG with minimal bilateral recording was employed (Bouilleret et al., 1999; Riban et al., 2002). Different chronic seizure onset patterns were evident in the same animal, and almost all seizures could be categorized as LVF or HYP. Importantly, seizure onset patterns resembled those in TLE (Velasco et al., 2000; Gnatkovsky et al., 2019) and following IHKA in rats (Bragin et al., 1999b; Bragin et al., 2005; Lévesque et al., 2012). Our results suggest that the prevalence of LVF or HYP seizures in an animal may change with time, a finding that is consistent with earlier observations in the rat pilocarpine- and IHKA-treated rats (Bragin et al., 1999b). This is important to be noted as it suggests changes in the seizure-generating network as epilepsy continues. On the other hand, characteristics of the LVF or HYP seizures did not significantly change. For example, both seizure types continued to be accompanied by stage 4-5 convulsions and were followed by prolonged postictal depression. On the other hand, HYP seizure duration increased with time. Therefore, the data presented here suggest that once initiated, HYP seizures may reverberate more with time, leading to a prolongation of the HYP seizure.

### D. High frequency oscillations (HFOs) occurred during seizures, at seizure onsets, as well as interictally

HFOs are considered hallmarks of epileptogenicity by numerous clinical (Jacobs et al., 2008; Weiss et al., 2016) and animal (Bragin et al., 1999a) studies and are currently used in the presurgical evaluation of patients with epilepsy (Zijlmans et al., 2019).

Although definitions and terms vary, we defined HFOs in a way that distinguished them from most normal oscillations that have high frequency, such as those between 100 and 200 Hz. Our definition was >250 Hz. We also used spectrograms to ensure peak frequency was indeed above 250 Hz. Note that others have used the term “pathological HFO (pHFO)” to refer to HFOs in epilepsy, but we have avoided that term because some HFOs occur in normal tissue (Engel et al., 2009; Pearce et al., 2014).

HFOs in our study were present at seizure onsets and during seizures, which has been reported in acute and chronic IHKA-induced seizures in the rat (Bragin et al., 1999b; Lévesque et al., 2012; Li et al., 2018) and in TLE (Weiss et al., 2016). We found frequent HFOs during periods of slow wave activity similar to reports of HFOs during delta activity in humans (Staba et al., 2004) and rats (Bragin et al., 1999a; Bragin et al., 2016). Importantly the NREM stage of sleep where delta oscillations are prominent appears to be the most useful to identify the seizure onset zone in TLE (Klimes et al., 2019). We also report HFOs outside the IHKA injection site, which suggests a wider epileptogenic network involving adjacent cortical and remote hippocampal areas (Sheybani et al., 2018).

There is currently only one mouse IHKA study that reported HFOs, but it mostly focused on power (Häussler et al., 2012) rather than the identification of individual HFO events during sleep, which is standard for the detection and evaluation of HFOs in clinical practice (Frauscher et al., 2017). Importantly, we found that both male and female mice showed HFOs. This is important because both sexes have not been studied before. These data suggest that HFOs, an abnormality with an important translational value, are in both sexes. The findings support the increasing need to expand the bandwidth normally used for preclinical studies to capture high frequency abnormalities in the EEG of animals with epilepsy.

### E. Hippocampal pathology at the IHKA injection site was consistent with Mesial Temporal Sclerosis (MTS)

Human tissue from many patients with intractable TLE shows extensive pyramidal cell loss. The pattern called MTS is characterized by neuronal death primarily in the hilus, CA3 and CA1 (Houser, 1999; Scharfman, 2006; Blümcke et al., 2012; Thom, 2014). In addition, the dentate gyrus is characterized by granule cell dispersion (Houser, 1990; Suzuki et al., 1995; Bouilleret et al., 1999; Riban et al., 2002). The original studies of IHKA showed extensive pyramidal cell loss and granule cell dispersion near the site of IHKA injection, with little evidence of neuropathology in the contralateral hippocampus (Suzuki et al., 1995; Bouilleret et al., 1999). In the animals that we studied after IHKA, we also saw extensive pyramidal cell loss near the site of IHKA injection, although there was variability. Quantification showed a significant reduction in pyramidal cells near the IHKA injection site in both CA3 and CA1, like MTS. We also observed significant granule cell dispersion in most animals, and when quantified, granule cell dispersion was reflected by an increase in granule cell area and thickness (post-IHKA compared to Saline).

The mechanisms by which KA induces cell death in pyramidal layers are thought to be excitotoxicity mediated by KA receptors, and due to prolonged seizures (Ben-Ari et al., 1980). To address a possible correlation between histopathology and seizures during SE, we plotted the number of stage 5 seizures during IHKA-induced SE and histopathological measurements (pyramidal cell length, GCL area, and GCL average thickness). We also plotted the relation between chronic seizures and histopathological measurements. The data showed a significant correlation between neuronal loss in the pyramidal cell layers and the number of stage 5 seizures during SE. The results are similar to those of the intraamygdala KA model in rats (Ben-Ari et al., 1980; Henshall et al., 2000) and mice (Araki et al., 2002) although in prior studies the investigators used other methods to quantify neuronal damage than the ones we implemented. Together these data support the idea that severe SE is needed for the neuronal loss in the pyramidal cell layers (Dingledine et al., 2014). This is notable because a long-standing assumption is that chronic epilepsy is more likely when there is greater neuronal loss (Dingledine et al., 2014).

It is important to note that we evaluated pyramidal cell loss at the end of our experiments, typically 12 wks post-IHKA. Others typically examine neuronal loss in the days following SE (Ben-Ari et al., 1980; Henshall et al., 2000; Araki et al., 2002).

However, we did assess some animals at early times and the neuronal loss in the pyramidal cell layer was similar. Moreover, in SE models the vast majority of neuronal loss occurs within 10 days of SE so we do not think that examining mice after 12 wks was a major limitation.

In contrast to the length of the pyramidal cell layer, measurements of granule cell dispersion were not correlated with acute or chronic seizures. This lack of correlation in our data might be attributed to the fact that granule cell dispersion is an alteration which is not dependent on SE severity or chronic seizures. Alternatively, the measures of SE or chronic seizures may not have been those that are related to granule cell dispersion. For example, we did not measure the power or duration of SE but the number of stage 5 seizures during SE because the animals were not vEEG monitored during SE. Power and duration of SE may be more sensitive as a measure of SE severity than the number of individual convulsive seizures. The same could be true of chronic seizures. On the other hand, the mechanisms underlying granule cell dispersion may be initiated by the way KA receptors modify the granule cells, independent of SE convulsive seizures or chronic epilepsy. Other possibilities also exist, based on proposed mechanisms for granule cell dispersion (Haas et al., 2002).

### F. Using IHKA to induce robust chronic epilepsy in mice

Our data suggest that the IHKA model in mice induces robust chronic epilepsy, unlike several past reports where chronic convulsive seizures are not noted, or the EEG evidence does not clearly reflect robust seizures (Kiasalari et al., 2016; Zhu et al., 2016; Runtz et al., 2018; Bielefeld et al., 2019; Li et al., 2020). Notably, some investigators have stated that they see chronic epilepsy after IHKA, but the evidence is not always presented (Kim et al., 2018; Sandau et al., 2019; Lai et al., 2020).

Why would our methods lead to more convulsive seizures after IHKA-SE than other methods? One potential explanation is related to the fact that our animals sustained IHKA-SE with multiple stage 5 seizures. In some other studies of IHKA, non-convulsive SE was reported (Bouilleret et al., 1999; Riban et al., 2002; Arabadzisz et al., 2005; Maroso et al., 2011). If it is true that there is more neuronal loss when there are more convulsive seizures during SE, and furthermore if it is true that the greater neuronal loss is, the more robust the chronic seizures, then our initiation of IHKA-SE with many stage 5 seizures could have led to more robust epilepsy.

Another possibility is that we did not implant animals with electrodes or cannula prior to SE (or immediately after IHKA injection), which is notable because in the past we have found that implanted animals have a higher seizure threshold ((Jain et al., 2019) and HES, unpublished).

We also handled mice more than is typical. Thus, we handled mice once or twice per day for the 2 days before IHKA injection and a longer time period afterwards to account for the behavioral stress related to the lack of social housing (Bernard, 2019; Manouze et al., 2019). Animals also were housed with miniature enclosures so that animals could enter the enclosure and not be seen. This type of environment would be likely to lower behavioral stress even more. A study in the pilocarpine model suggested that single housing may increase seizures by a factor of 16 (Manouze et al., 2019).

Single housing might thus explain why very frequent and brief (3-7 sec) epileptiform abnormalities have been reported in IHKA-treated mice in the past (Kim et al., 2018; Sandau et al., 2019; Lai et al., 2020; Rusina et al., 2021). It would be possible that our approach that included frequent handling combined with enriched housing reduced these very frequent abnormalities thereby allowing the expression of robust seizures. These ideas are consistent with past studies suggesting that stress plays a major role in seizure induction and in chronic epilepsy in both animals (MacKenzie and Maguire, 2015) and humans (Lang et al., 2018), although the relationship between stress and seizures/epilepsy is complex (Gunn and Baram, 2017).

Another factor is that we modified the pH of the solution of KA we injected so it was closer to a physiological pH. In the past, the pH may not have been monitored, and if not, a very acidic solution would have been injected because the pH of our solution, before adding a base to bring it closer to the physiological range, was approximately 4.0. In the past, we found that bringing the solution of pilocarpine to pH 7.4 before injection played a very large role in the consequences of pilocarpine with fewer seizures when pH was not controlled (Scharfman et al., unpublished). Finally, we found that seizures can be robust, but if a time when seizures are limited is the only time that is studied, one might get a false sense of few seizures when animals actually had robust epilepsy at other times.

### G. Limitations of the study

One of the limitations of our study is that a subset of the experiments used some mice that were transgenic. However, the mice were designed to express Cre recombinase conditionally, and only in small groups of neurons. Since we did not conditionally activate Cre, we assume the mice that were hemizygous acted like the background strain, C57BL/6. In addition, wild type mice were also used. In fact, we found no evidence that the hemizygous and wild type mice differed in acute effects of SE or chronic seizures as shown in **Table 2**. We also used C57BL6 mice from 2 different vendors and results were similar (**Table 2**). Therefore, we do not think the use of transgenic mice was a major limitation.

We also did not assess the EEG during SE, because all mice exhibited robust convulsive SE. Also, the presence of cannulas and electrodes could have influenced SE.

## ACKNOWLEDGEMENTS

We thank Drs. Annamaria Vezzani, Esther Krook-Magnuson, Ivan Soltesz, and Peter West for sharing their methods for injecting KA in mice. We also thank members of the Scharfman lab for fruitful discussions and John LaFrancois for sharing his method for intrahippocampal injection in mice. This work was supported by the National Institutes of Health [grant number NIH R01 NS106983] and the New York State Office of Mental Health.

## CONTRIBUTION OF AUTHORS

**Christos P. Lisgaras and Helen E. Scharfman**: Conceptualization, **Christos P. Lisgaras**: Data curation, **Christos P. Lisgaras and Helen E. Scharfman**: Formal analysis; **Helen E. Scharfman**: Funding acquisition, **Christos P. Lisgaras**: Investigation; **Christos P. Lisgaras**: Methodology, **Helen E. Scharfman**: Project administration, **Helen E. Scharfman**: Resources, **Christos P. Lisgaras**: Software, **Helen E. Scharfman**: Supervision, **Christos Panagiotis Lisgaras**: Validation, **Christos P. Lisgaras and Helen E. Scharfman**: Visualization, **Christos P. Lisgaras**: Roles/Writing - original draft, **Helen E. Scharfman and Christos P. Lisgaras**: Writing - review & editing.

## Abbreviations

IHKA: Intrahippocampal kainic acid
KA: Kainic acid
TLE: Temporal lobe epilepsy
LVF: Low-voltage fast
HYP: Hypersynchronous
HFOs: High frequency oscillations
SE: Status Epilepticus
vEEG: video electroencephalography

## SUPPLEMENTAL MATERIAL

**Supplemental Table 1.** Data for Saline-injected controls. Data for Saline-injected controls are shown. No spontaneous seizures and no HFOs were detected. RFC: right frontal cortex; LFC: left frontal cortex; LHPC: left hippocampus; RHPC: right hippocampus.

| Cage ID | Sex | Genotype | Experimental use | Treatment | Dose (nL) | Timepoint of EEG recording start (wks) | # HFOs RFC | # HFOs LFC | # HFOs LHPC | # HFOs RHPC | # Seizures |
| --- | --- | --- | --- | --- | --- | --- | --- | --- | --- | --- | --- |
| 99-1 | Female | Cre -/- | EEG | Saline | 70 | 10 | 0 | 0 | 0 | 0 | 0 |
| 99-3 | Female | Cre +/- | EEG | Saline | 100 | 10 | 0 | 0 | 0 | 0 | 0 |
| 99a-5 | Male | Cre +/- | EEG | Saline | 70 | 10 | 0 | 0 | 0 | 0 | 0 |
| 100-4 | Female | Cre -/- | EEG | Saline | 100 | 10 | 0 | 0 | 0 | 0 | 0 |
| Jax a-3 | Male | NA | EEG | Saline | 100 | 2 | 0 | 0 | 0 | 0 | 0 |
| Jax a-4 | Male | NA | EEG | Saline | 100 | 2 | 0 | 0 | 0 | 0 | 0 |

**Supplemental Figure 1.**
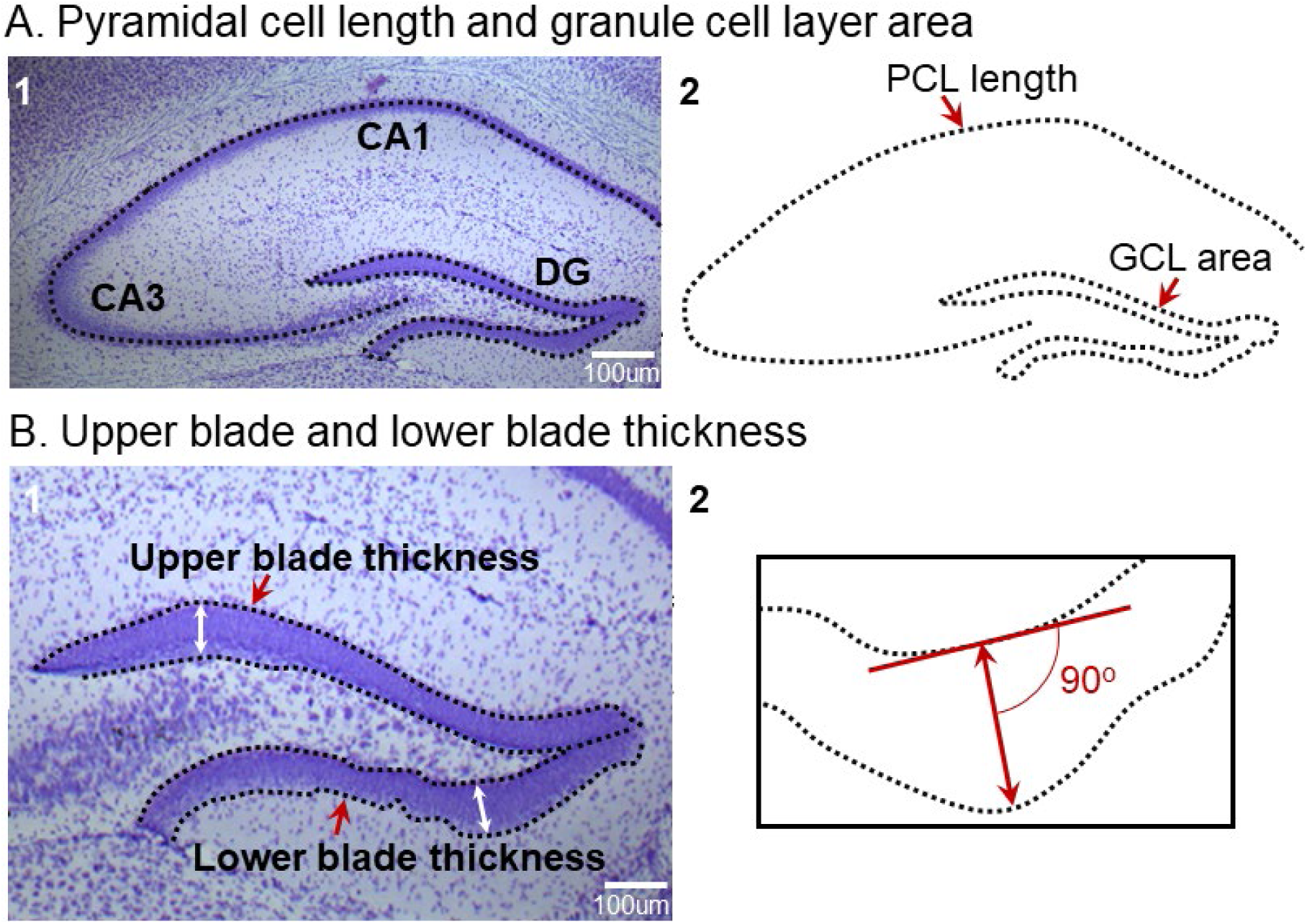
Quantification of hippocampal pathology 12 wks post-IHKA. Methods used for cresyl violet-stained sections from dorsal hippocampus near the area of the IHKA injection. Measurements were made using ImageJ as discussed in the Methods. **(A)** A cresyl violet-stained section (A1) and the same image without the stained section (A2) shows the lines used for measurements. For pyramidal cell length (PCL), a line was drawn using the freehand line tool from CA3c to the end of CA1. The line included those areas where pyramidal cells were contiguous, i.e., there was <1 soma width between cell bodies. The CA1 border with the subiculum and the CA3c border with the hilus was the end of CA1 and CA3c respectively. These areas were defined in the Methods. For the measurement of the granule cell layer (GCL) area, we traced the borders of the GCL using the freehand tool and calculated the area within the borders. The border of the GCL and hilus and border of the GCL and IML were defined by the point where adjacent cells became >1 GC soma width apart. **(B)** For GCL width, we used the straight-line tool to measure the thickest part of the upper and lower blade (red arrows; B1) based on the traced borders of GCL. The white line with arrows (B1) was perpendicular to the length of the GCL as shown in B2. We averaged 2 measurements for an estimate of upper blade width of a section and then did the same procedure for the lower blade to yield the estimate for the lower blade width of that section.

**Supplemental Figure 2.**
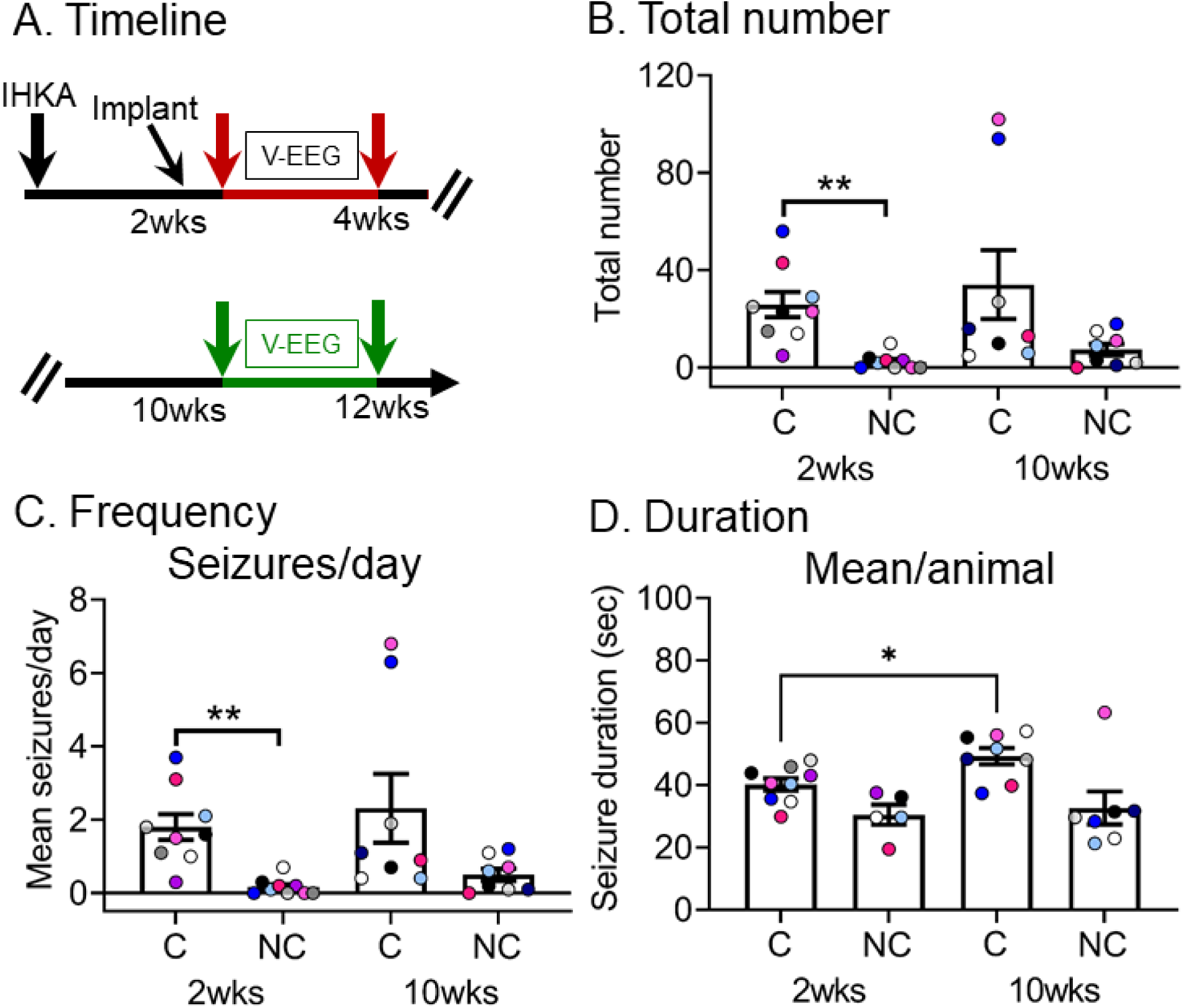
Quantification of convulsive and non-convulsive seizures for 2-4 and 10-12 wks post-IHKA for animals that recorded at both timepoints. **(A)** Experimental timeline shows the times when measurements were made; 2-4 wks (red arrows) and 10-12 wks (green arrows) post-IHKA. **(B)** The total number of chronic convulsive and non-convulsive seizures recorded during 2 wks of continuous vEEG is shown for the 2-4 wk timepoint (n=9 animals; 6 males; 3 females) and 10-12 wk timepoint (n=8 animals; 6 males, 2 females). These animals were the same except for one that was removed from the study and therefore could not be included in the 10-12 wk timepoint. The number of convulsive seizures was significantly greater than the number of non-convulsive seizures (paired t-test, *t*_crit_ = 4.16, p=0.030), suggesting that the majority of seizures were convulsive. **(C)** Same as B, but seizure frequency is shown. The frequency of convulsive seizures was significantly greater than non-convulsive seizures (Wilcoxon signed rank test, p=0.03), again suggesting that there were more convulsive seizures in the animals. **(D)** Same as C, but seizure duration is shown. Seizure duration was calculated as an average per animal. Convulsive seizures lasted longer at 10-12 wks vs. 2-4 wks (paired t-test, *t*_crit_ = 3.14, p=0.02). Non-convulsive seizures did not differ between timepoints (Wilcoxon signed rank test, p>0.05).

**Supplemental Figure 3.**
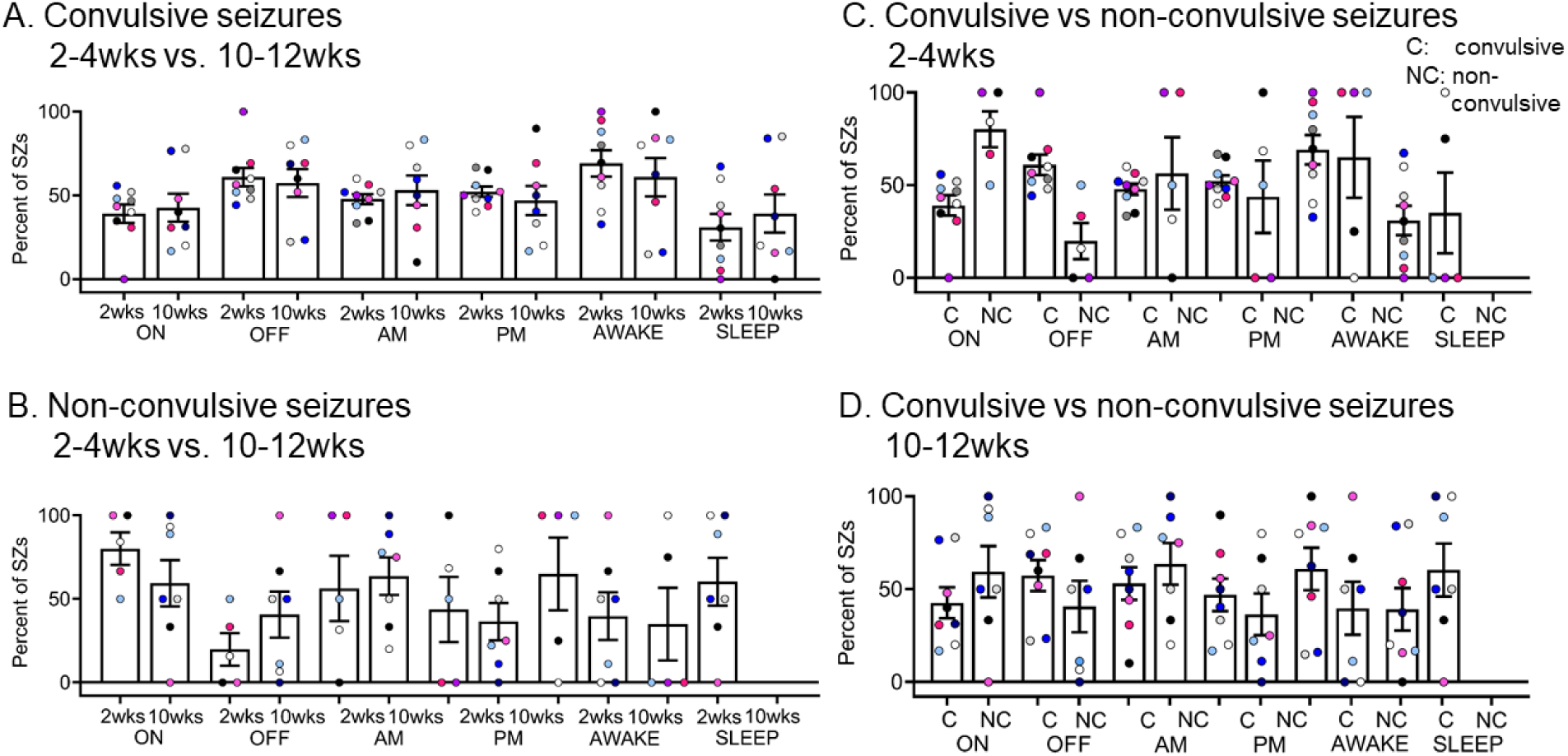
Percent of convulsive and non-convulsive seizures at the two time points (2-4, 10-12 wks): effects of light, time of day and sleep. **(A)** Percent of convulsive seizures that occurred when lights were on or off, a.m. or p.m., and awake or sleep for the two time points (2-4 and 10-12 wks). Only animals recorded at both timepoints are included. **(B)** Same as in A but for non-convulsive seizures. **(C)** Data were compared using just the 2-4 wk timepoint. **(D)** Same as C, but for the 10-12 wk timepoint. There were no significant differences (Wilcoxon signed rank test, p>0.05) most likely due to the variability from one animal to the next.

**Supplemental Figure 4.**
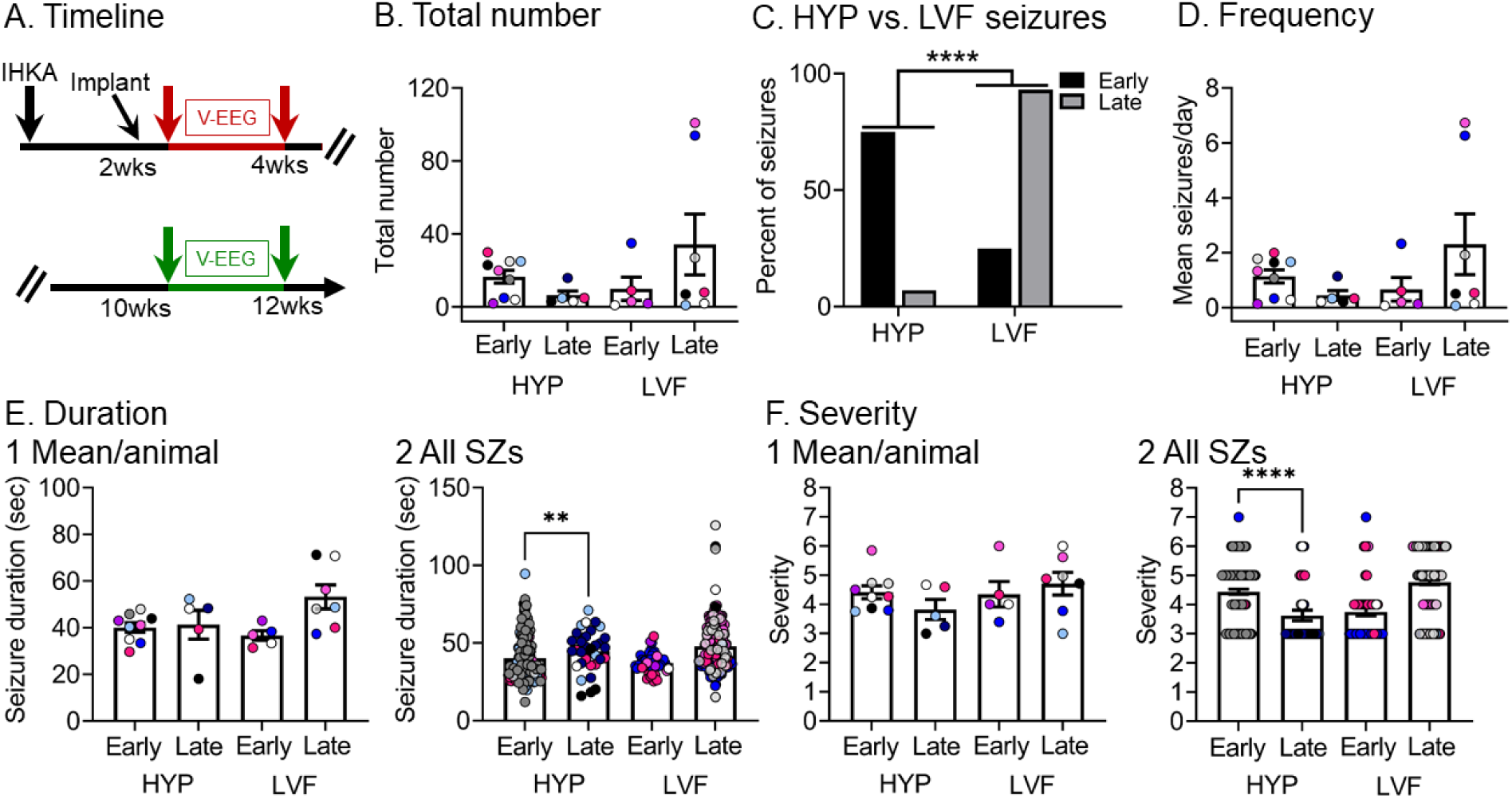
Seizure onset patterns at 2-4 and 10-12 wks for convulsive seizures only. **(A)** Experimental timeline of the study. Data from the early (red arrows) and late (green arrows) timepoint are included in this figure. There were 9 mice at 2-4 wks (7 males, 2 females). The same mice were recorded at 10-12 wks except for two of the males that could not be recorded at 10-12 wks. **(B)** The total number of chronic convulsive HYP and LVF seizures recorded during 2-4 wks and 10-12 wks. No significant differences were found for HYP or LVF seizures between the 2 timepoints (HYP: Wilcoxon signed rank test, p=0.12, LVF: Wilcoxon signed rank test, p=0.37). **(C)** Percent of HYP and LVF seizures at the 2-4 and 10-12 wk timepoints. The percent of HYP vs. LVF seizures is different between timepoints with HYP seizures dominating the earlier timepoint and LVF seizures the late timepoint (Fisher’s exact test, p<0.0001). **(D)** Same as B, but convulsive seizure frequency is shown. Neither the frequency of HYP nor LVF seizures changed significantly between timepoints (Wilcoxon signed rank tests, p=0.12). **(E)** Same as B, but convulsive seizure duration is plotted. Seizure duration was calculated as the average per animal in E1 and for all seizures in E2. There were no significant differences for HYP and LVF seizures between the 2 timepoints. when seizure duration was calculated as a mean per animal (HYP: Wilcoxon signed rank test, p=0.87, LVF: Wilcoxon signed rank test, p=0.25). However, overall, HYP seizures lasted significantly longer at the late timepoint when all seizures were compared (Wilcoxon signed rank test, p=0.002; E2). **(F)** Same as B, but convulsive seizure severity is shown. Convulsive seizure severity was calculated as the average per animal in F1 and for all seizures in F2. There were no significant differences when seizure severity was calculated as a mean per animal (Wilcoxon signed rank tests, p>0.05; F1). However, HYP seizures were less severe at the late timepoint when all seizures were compared (Wilcoxon signed rank test, p<0.0001; F2) possibly because of the larger sample size.

**Supplemental Figure 5.**
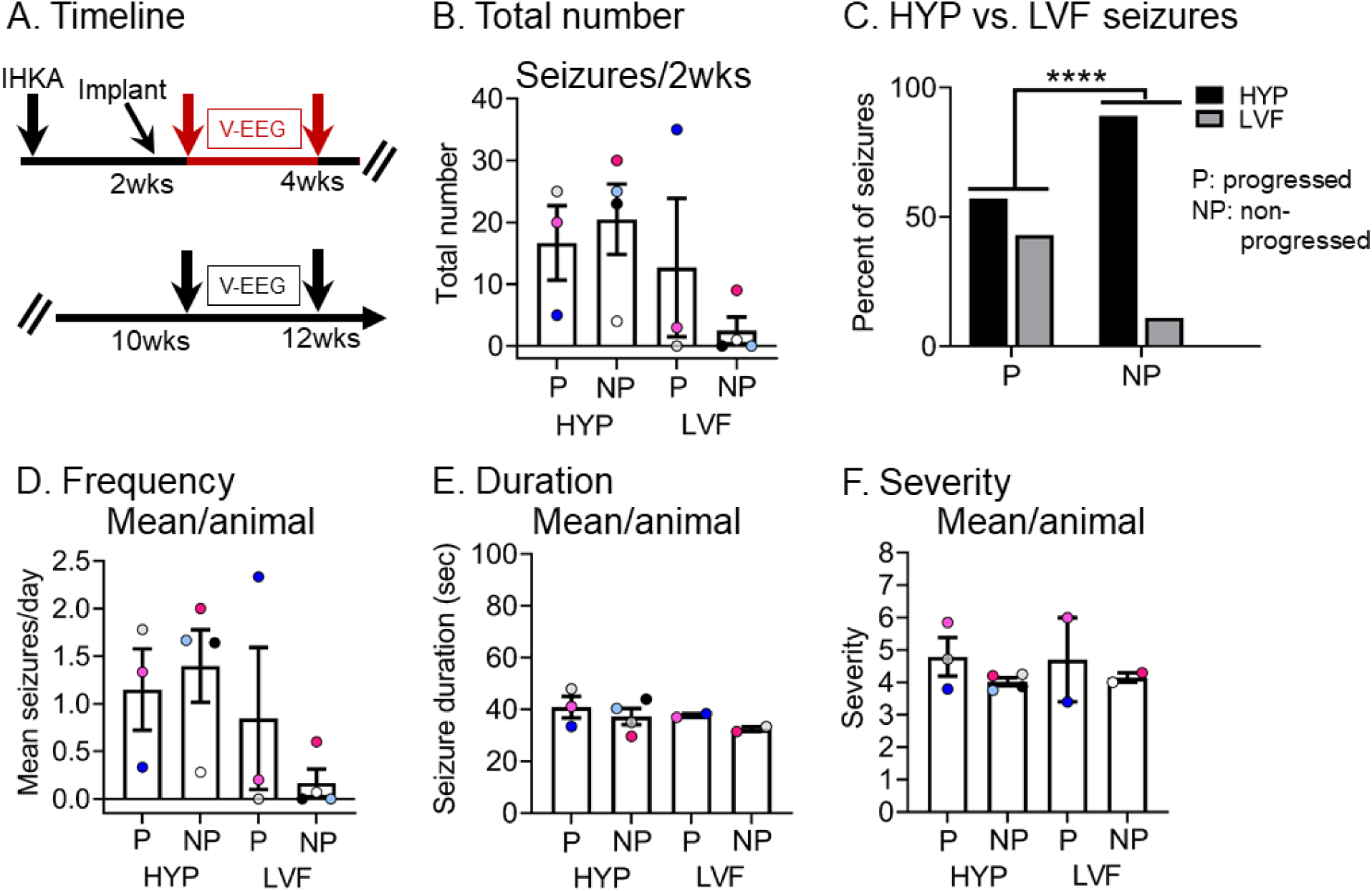
Comparison of different convulsive seizure onset patterns in mice that did or did not show increased seizures from 2-4 to 10-12 wks. **(A)** Experimental timeline of the study as shown before. Data from the early timepoint (red arrows) are used for this figure. The comparisons are between mice with seizures that were most frequent at 2-4 wks and mice that with seizures that were most frequent at 10-12 wks. There were 2 mice/group. For simplicity, worsening at 10-12 weeks is termed progressed (P) and when worsening did not occur, the term non-progressed (NP) is used. However, these terms should be used cautiously because only two timepoints were compared. **(B)** The total number of convulsive HYP and LVF seizures recorded during 2-4 wks is shown for two subsets of mice, those with seizures that increased between 2-4 wks and 10-12 wks (progressed, P) and those mice with seizures that decreased (non-progressed, NP). There were 7 mice (5 males, blue; 2 females, pink). **(C)** The percentages of HYP or LVF seizures at 2-4 wks were compared between P and NP mice. Mice that showed progression (P) had a significantly different proportion of HYP seizures relative to LVF seizures depending on their progression (P) or non-progression (NP; Fisher’s exact test, p<0.0001). The data support the view that HYP seizures dominated non-progressed mice in contrast to progressed mice where both patterns seem to be equally prevalent. **(D)** Same as B, but seizure frequency is shown. **(E)** Same as B, but seizure duration is plotted. Seizure duration was calculated as an average per animal. **(F)** Same as B, but seizure severity is shown. Convulsive seizure severity was calculated as an average per animal.

**Supplemental Figure 6.**
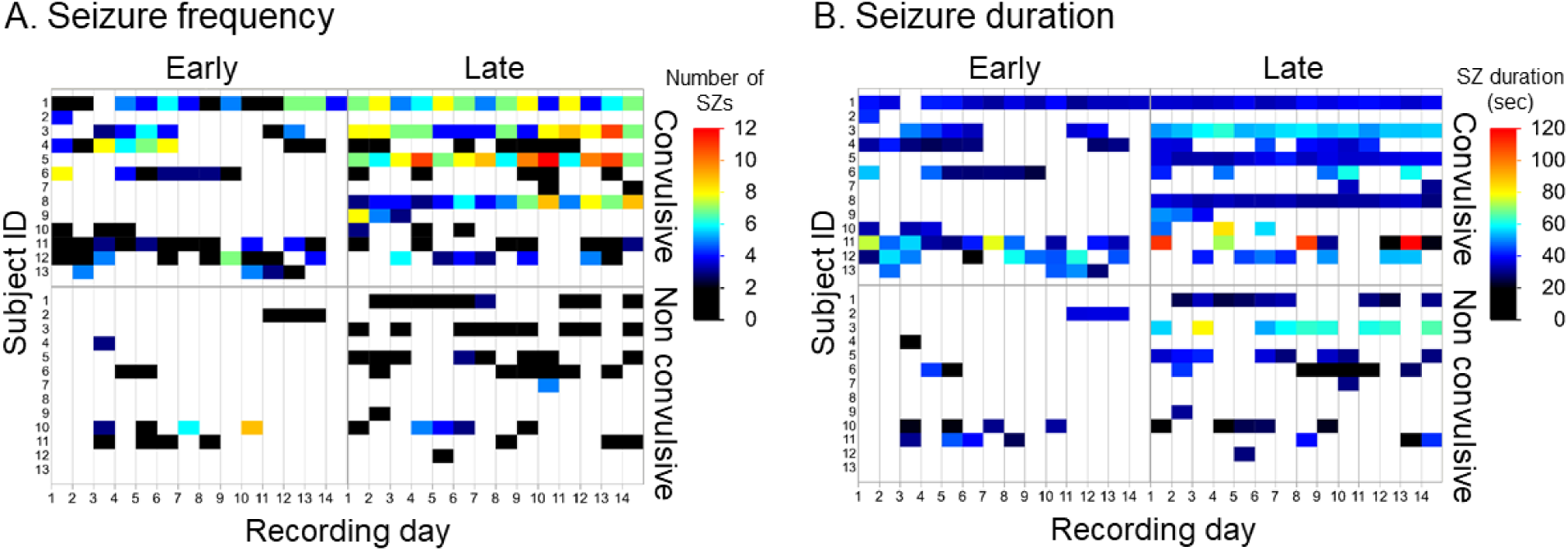
Seizure frequency and duration of convulsive and non-convulsive seizures for each individual animal at each timepoint. **(A)** Seizure frequency for each animal included in the study. Upper panel shows convulsive seizures and lower panel non-convulsive seizures. Left part of the panel shows the early time point and right part the late timepoint. Color scale showing the number of seizures is shown on the right. Warmer colors indicate a greater number of seizures. **(B)** Seizure duration for each animal included in the study. Upper panel shows convulsive seizures and lower panel non-convulsive seizures. The left half of the plot shows the early time point and the right part of the plot shows the late timepoint. The color scale showing the duration of seizures is shown on the right. Warmer colors indicate longer seizures.

**Supplemental Figure 7.**
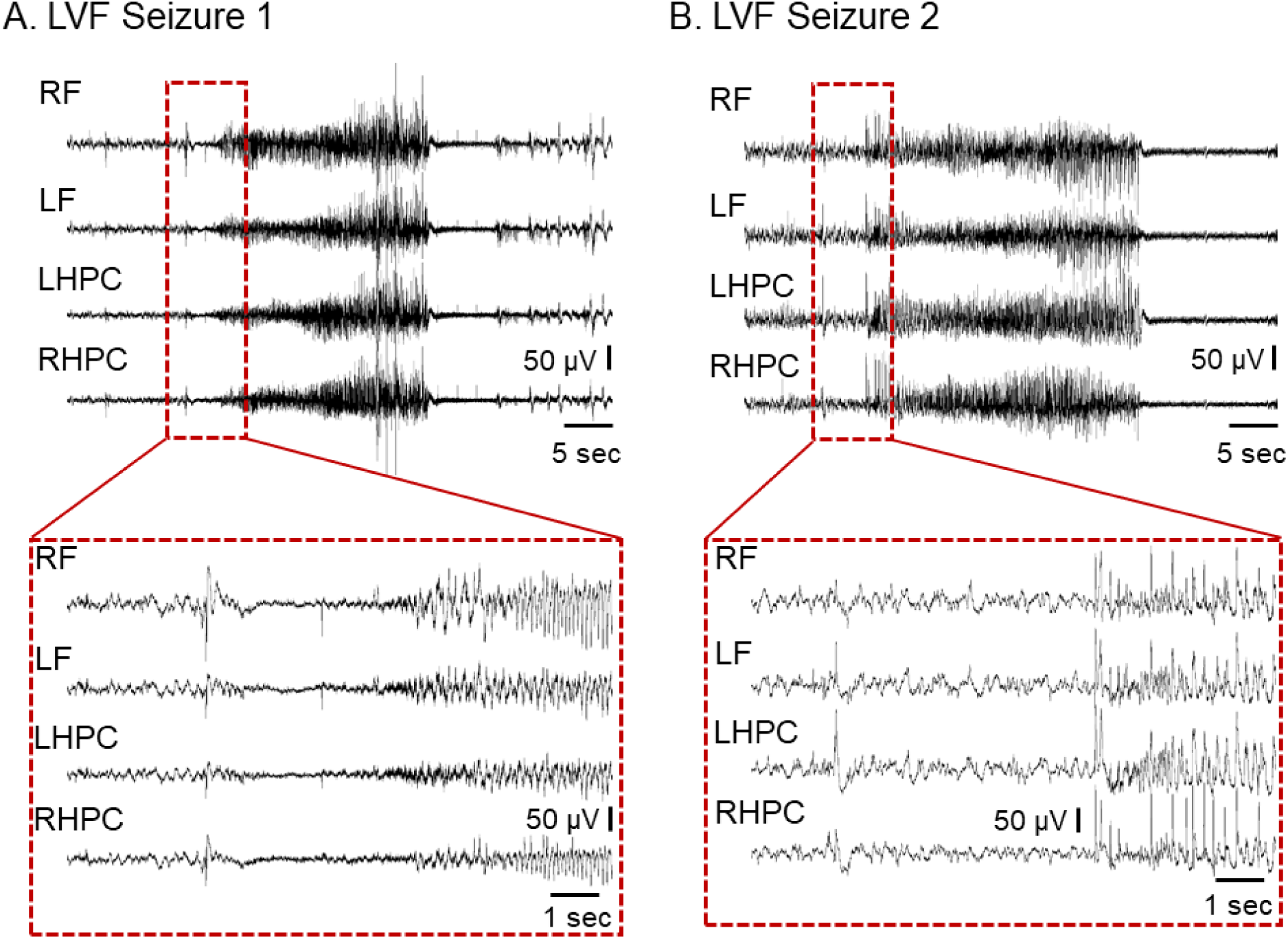
Additional examples of LVF seizures. The start of the seizure outlined by the red box at the top is explanded below. Abbreviations are the same as the main text. **(A)** A LVF seizure from animal #5. Note the prominent “sentinel” spike evident in all four leads, occurring almost at the same time. Afterwards is a brief suppression of background EEG, also in all four leads. Subsequently the fast rhythmic activity begins and becomes large in amplitude. The seizure terminates with a sudden cessation of this large activity. **(B)** A LVF sezure from animal #12. Note the similarity to A, but the suppression of the background EEG after the sentinel spike is less clear.

**Supplemental Figure 8.**
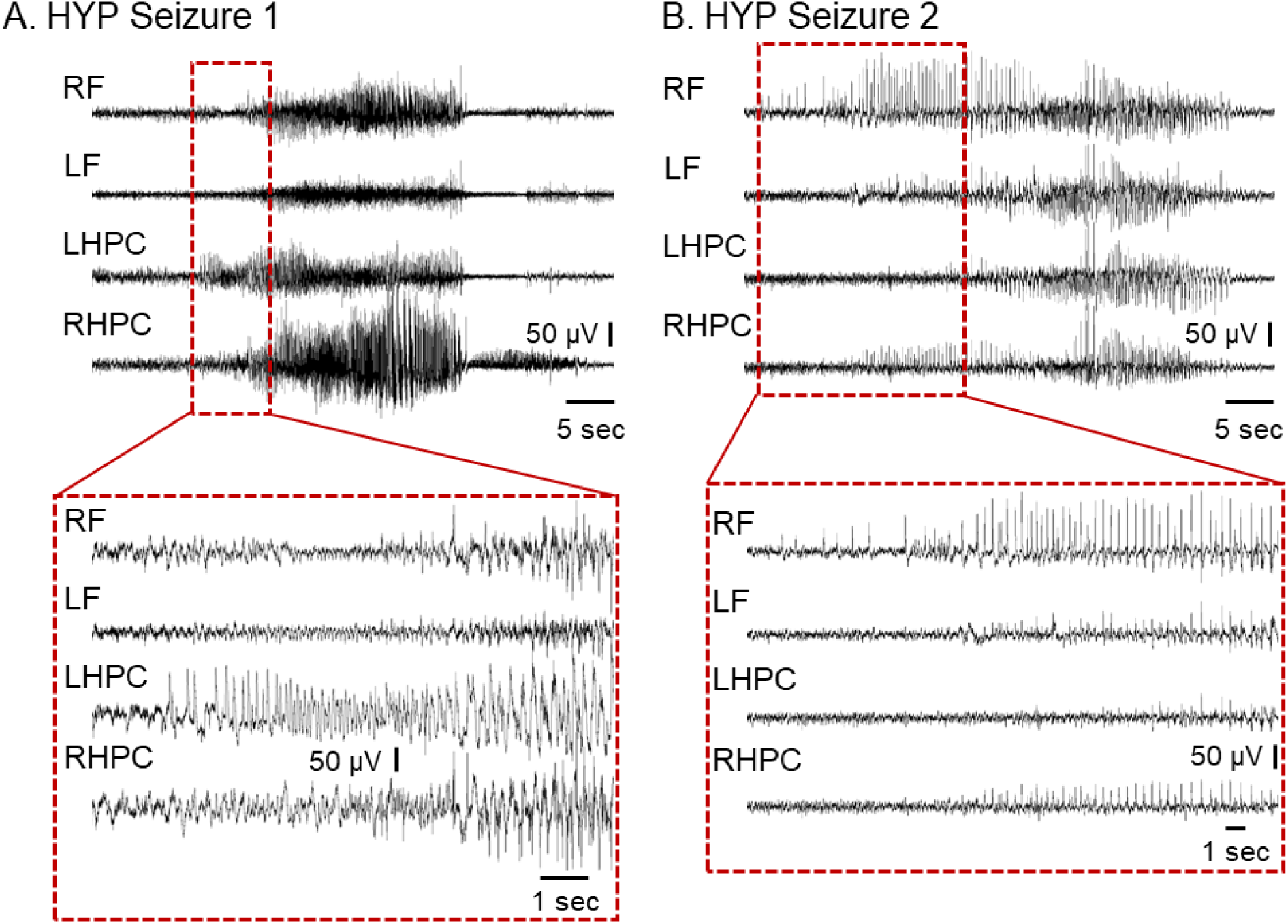
Additional examples of HYP seizures. The start of the seizure outlined by the red box at the top is explanded below. Abbreviations are the same as the main text. **(A)** A HYP seizure from animal #4. Note the absence of a sentinel spike. Instead seizures start with fast rhythmic activity that increases in amplitude and is complex. The frequencies of the activity are highly variable. Also, the seizure does not start simultaneously in all four leads. **(B)** A HYP sezure from animal #9.

